# Rapid and Complete Inactivation of Enveloped Viruses by Electrochemical Disinfection: Unraveling the Contribution of Reactive Chlorine Species

**DOI:** 10.1101/2025.11.14.688411

**Authors:** Chenyang Xu, Chengxu Jiang, Shih-Han Sun, Peiyang Sang, Ruipeng Shao, Xiaohong Zhou, Xiaoyuan Zhang, Yancheng Liu, Xianghua Wen, Xia Huang

## Abstract

Waterborne viruses pose significant threats to global water safety. Electrochemical (EC) disinfection is considered a promising next-generation technology to address this challenge. However, its efficacy against enveloped viruses remains inadequately under-stood. In this study, we employed bacteriophage Phi6 as a surrogate to systematically eval-uate the inactivation efficiency and damage mechanisms of enveloped viruses in a flow-through EC reactor. Experimental results indicated that Phi6 was more susceptible to EC disinfection than common bacterial surrogates. Comprehensive damage was observed across all major viral components, including structural proteins, the lipid envelope, and the RNA genome. The electrogenerated reactive chlorine species (RCS) were identified as the primary agent responsible for this inactivation, exhibiting higher reactivity than conven-tional free chlorine. Furthermore, radical chlorine species were also confirmed to be pro-duced during EC disinfection, contributing to rapid and extensive viral inactivation. This study demonstrates the feasibility of EC disinfection for inactivating enveloped viruses and provides deeper mechanistic insights into its highly efficient viral inactivation performance.

**SYNOPSIS:** Our results elucidate the rapid inactivation and thorough damage of the enveloped virus surrogate Phi6 by electrogenerated reactive chlorine species during electrochemical disinfection.

**TOC:** 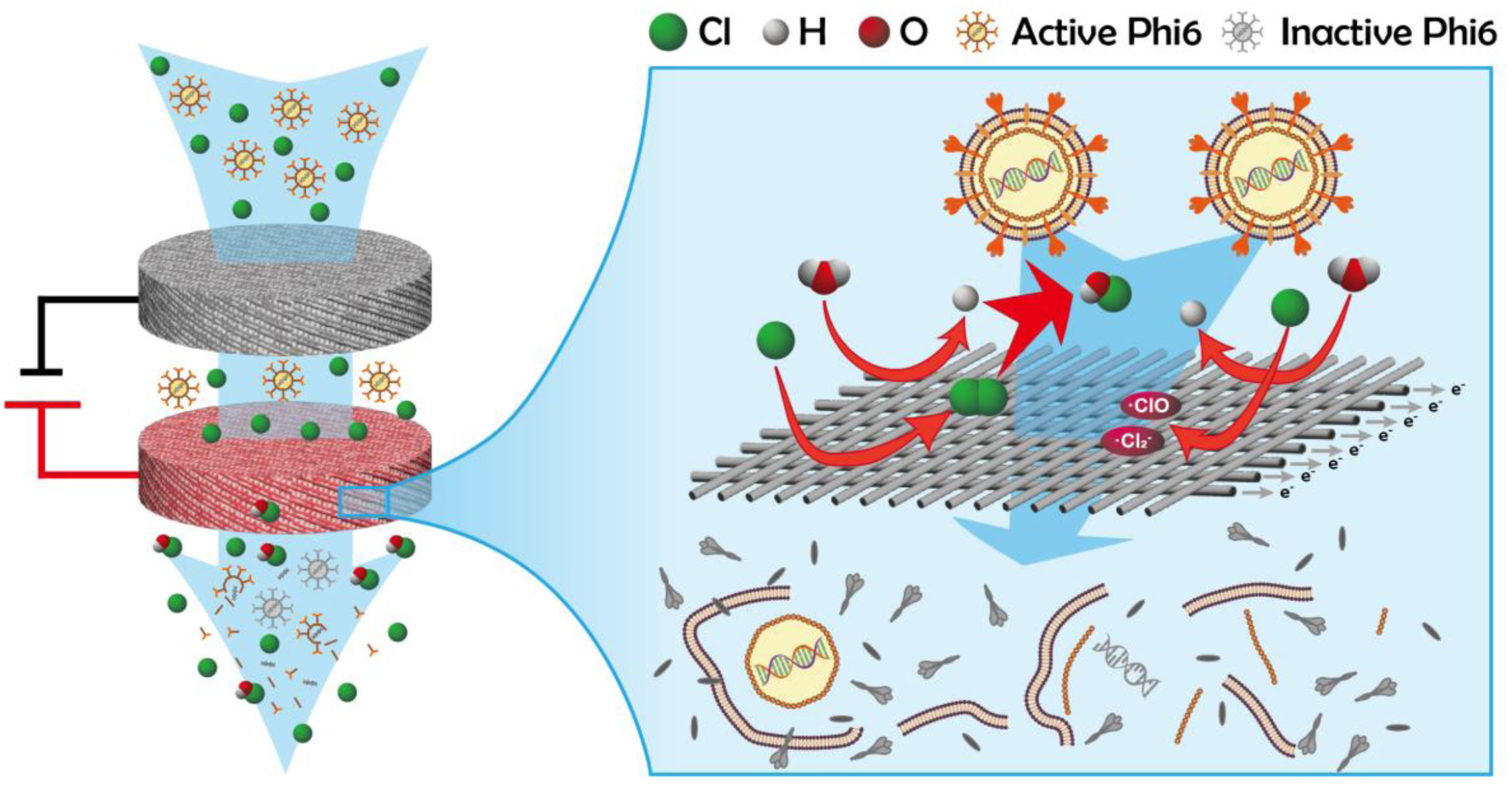

## 1 INTRODUCTION

Human pathogenic viruses are a group of small but dangerous pathogens, imposing a significant burden on global public health^1,2^. To date, extensive studies have sufficiently demonstrated that a wide variety of viruses can be released into domestic wastewater through the excreta of infected individuals^1,3,4^. Moreover, substantial evidence indicates that some viruses may maintain their infectivity for several days or even longer periods in raw sewage and other water bodies, which underscores the potential of wastewater and reclaimed water as covert pathways for viral transmission^5,6^. Traditionally, research and control strategies targeting viral transmission in wastewater have predominantly centered on non-enveloped viruses, such as noroviruses^7,8^, enteroviruses^9,10^, and adenoviruses^8^. In recent years, however, the outbreaks and spread of emerging enveloped viruses, including Ebola virus, severe acute respiratory syndrome coronavirus 2 (SARS-CoV-2), monkeypox, and highly pathogenic avian influenza, have brought the potential transmission risks and control of enveloped viruses in wastewater to the forefront of attention^1,11,12^. Though some studies suggested that enveloped viruses may exhibit relatively poor persistence in the environment, their extremely low infectious doses and high pathogenicity still warrant careful consideration for water safety^13,14^. Meanwhile, current investigations have indicated that conventional wastewater treatment processes cannot completely remove or inactivate all human-infectious viruses^1,3^. Therefore, it is crucial to develop more effective and dependable virus inactivation technologies to ensure urban water safety.

Electrochemical (EC) technology, characterized by its high energy efficiency, ease of operation, and minimal chemical addition, has emerged as a promising candidate for next-generation disinfection approaches^15,16^. Under EC treatment, multiple reactions involving electric field effect^17^, direct anode oxidation^18,19^, indirect anode oxidation based on reactive oxygen species (ROS, including hydroxyl radical, singlet oxygen, i.e.) ^18,19^ or reactive chlorine species (RCS, including free chlorine, chlorine radical, dichlorine radical, chlorine oxide radical, i.e.) ^17,20^, and cathodic EC processes^21^ have been proven to inactivate pathogens effectively. To date, considerable efforts have been devoted to investigating and optimizing microbial inactivation by EC disinfection. However, the vast majority of this research has focused on bacteria, with viruses receiving relatively little attention. Among those studies addressing viruses, non-enveloped viruses—notably bacteriophage MS2^22,23^, feline calicivirus (FCV)^24^, and recombinant adenovirus (rAdeV)^25^—are frequently employed as surrogates. In contrast, studies focusing on the inactivation of enveloped viruses under EC treatment remain particularly scarce. Therefore, systematic evaluation is essential to determine whether EC disinfection could effectively inactivate enveloped viruses.

On the other hand, most previous studies have merely focused on evaluating the loss of viral infectivity by EC treatment, while the mechanisms underlying viral damage through the EC disinfection have garnered considerably less attention^14,22^. Relevant studies have demonstrated that viruses often exhibit distinct patterns of damage when inactivated by various methods. For instance, reaction with chlorine dioxide (ClO_2_) causes protein damage^26^, inactivation by free chlorine (FC) compromises viral structural integrity^22^, and exposure to ultraviolet (UV) irradiation inflicts genomic injury^23^. Inspired by these findings, it is essential to elucidate the damage mechanisms of viruses within the EC disinfection systems and identify the specific electrode reactions responsible for such damage. These results will not only advance our understanding of the underlying mechanisms of EC dis-infection but also offer valuable insights for the further optimization and enhancement of EC disinfection technologies.

In this study, the bacteriophage Phi6 was employed as a model surrogate to investigate the inactivation efficacy and underlying mechanisms of enveloped viruses during the EC disinfection^27^. The reliable inactivation performance was quantified in a representative flow-through EC reactor and benchmarked against commonly used bacterial surrogates. Subsequently, comprehensive molecular-biology analyses were applied to systematically assess the multi-level damage inflicted upon viral proteins, membranes, structural integrity, and RNA genomes. A complementary suite of controlled experiments was further con-ducted to pinpoint the principal electrochemical reaction responsible for viral inactivation. Our work elucidates the inactivation rate and the damage profile imposed by EC reactions on enveloped viruses and conclusively reveals the primary role of electrogenerated RCS in viral inactivation.

## 2 MATERIALS AND METHODS

### 2.1 Phi6 Cultivation and Purification

Bacteriophage Phi6 and its host *Pseudomonas syringae* (*P. syringae*) were purchased from ATCC. The virus stock was prepared as previously described with some modifications^28^. Briefly, *P. syringae* was propagated in tryptone soybean broth (TSB) at 26 ± 2 °C to an optical density (OD) of 0.20 ± 0.10. Phi6 stoke was then inoculated into the bacterial culture to achieve a multiplicity of infection (MOI) of approximately 0.1. The culture was incubated under the same conditions until the solution became completely transparent with visible debris. To harvest the phages, the lysate was centrifuged at 8,000 × g for 15 min and then filtered through 0.45 μm syringe filters. Subsequently, the resulting Phi6 solution was purified using the 30-kDa ultrafiltration centrifuge tubes (UFC703008, Millipore) and concentrated by approximately 100-fold. The final obtained Phi6 stock exhibited a phage titer of ∼10^11^ PFU mL^-1^, with the total organic carbon (TOC) below 1,000 mg L^-1^, ensuring that the carried-over organic matter would not significantly interfere with subsequent EC disinfection after a 1,000-fold dilution. The final stocks were aliquoted and stored at -80 °C until use to avoid repeated freeze-thaw cycles.

### 2.2 Electrochemical inactivation tests

The inactivation tests were conducted in a typical EC reactor with carbon fiber felt (CFF) electrodes, as shown in **Figure S2**. The fundamental physical and chemical properties of CFF electrodes were represented in **Table S1**. All the CFF electrodes were sequentially washed three times with acetone, ethanol, and ultrapure water to remove any potential contaminants. The influent solution was prepared by diluting the Phi6 stock 1000-fold in a 10 mM NaCl solution, yielding an initial titer of >10^7^ PFU mL^-1^. The virus-spiked solution was pumped into the reactor from the cathode to the anode at a flow rate of 5 mL min^-1^. The hydraulic retention times (HRT) for the entire reactor and the anode compartment were approximately 300 s and 60 s, respectively. For each experiment, the reactor was initially operated for 30 min to establish adsorption equilibrium before applying various cell voltages. All the effluent samples were collected at least 15 min after voltage initiation. Given that the pre-experiment has verified the absence of measurable residual disinfecting capacity in the effluent (**Figure S3**), chemical quenchers were omitted to avoid interference with disinfection efficacy assessment. The residual infectious titer of Phi6 was determined through the double-layer agar plaque assay. The inactivation efficiency was expressed as a log reduction (**Equation (**1)):

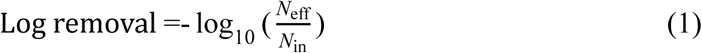

where *N*_in_ is the initial infectious titer before inactivation (PFU mL^-1^), *N*_eff_ is the re-sidual infectious titer after inactivation. Furthermore, the inactivation effects on two typical bacteria surrogates, *Escherichia coli* and *Bacillus subtilis* (*E. coli* and *B. subtilis*, details in **Text S2**), were also investigated under the same conditions for comparison. All the inactivation tests in different conditions were replicated at least twice.

### 2.3 Reverse Transcription Quantitative Polymerase Chain Reaction (RT-qPCR) and Propidium Monoazide (PMA)-mediated RT-qPCR Assays

The same samples collected for determining inactivation efficiency were also employed for the assessment of completeness and genomic injury. Briefly, total nucleic acids were extracted from 200 μL effluent samples using the Magnetic Viral DNA/RNA Kit (Tiangen, DP438) according to the manufacturer’s instructions. Subsequently, 5 μL of the eluted nucleic acid was utilized for the determination of genomic injury through RT-qPCR with previously reported primer/probe set^29^ and the FastKing One Step RT-qPCR Kit (Probe, Tiangen, FP314). Details on primer/probe sequences, the RT-qPCR reaction thermocycling protocol, and essential quality control (QC) information are provided in **Text S3**, **Table S2**, and the attached MIQE checklist. Furthermore, PMA-mediated RT-qPCR was employed to differentiate between intact and damaged viral particles following EC disinfection. The detailed protocols and verification tests for PMA staining and photo-inactivation are detailed in **Text S4** and **Table S3**.

### 2.4 Sodium Dodecyl Sulfate-Polyacrylamide Gel (SDS-PAGE) Electrophoresis, Transmission Electron Microscopy (TEM), and Fourier Transform Infrared Spectroscopy (FT-IR) Assays

Owing to the strict requirements of these molecular biology techniques for highly purified virus preparations to eliminate interference from residual bacterial debris, the viral stocks used for the SDS-PAGE, TEM, and FT-IR analysis were propagated using an optimized protocol and subjected to deep purification via ultracentrifugation until SDS-PAGE electrophoresis confirmed the absence of non-viral proteins (details in Text S5). For the EC disinfection, ∼10^12^ PFU of deeply-purified Phi6 were spiked into 500 mL of a pre-filtered 10 mM NaCl solution and treated in the EC reactor as described in Section 2.2. Following a 15-minute equilibration, the entire effluent was collected for subsequent concentration and characterization. Briefly, the effluent was first concentrated to 1 mL using 70 mL 10-kDa MWCO ultrafiltration centrifugal devices (UFC701008, Millipore), transferred to 2 mL 10-kDa MWCO centrifugal tubes (UFC201024, Millipore), and further reduced to a final volume of 100 µL. Aliquots were then processed as follows: (i) SDS-PAGE electrophoresis to assess protein damage; (ii) lyophilization, reconstitution in deuterium oxide (D₂O), and air-drying on an ATR crystal for FT-IR spectroscopy; (iii) deposition onto glow-discharged copper grids, negative staining with 2 % uranyl acetate, and TEM imaging. Detailed protocols are provided in Text S6.

### 2.5 EC-generated active species characterization

The electrogenerated FC and H_2_O^2^ were quantified by the N, N-diethyl-p-phenylene-diamine (DPD) colorimetric method and the potassium titanium oxalate method, respectively. A set of probe compounds, nitrobenzene (NB), benzoic acid (BA), 1,4-dimethoxybenzene (DMOB), and carbamazepine (CBZ) were chosen as the probe set to verify the existence of hydroxyl radical (·OH), chlorine radical (·Cl), dichlorine radicals (·Cl_2_-), and chlorine oxide radical (·ClO)^30,31^. Meanwhile, two broadly used scavengers, tert-butyl alcohol (TBA) and methallyl alcohol (MAA), were also applied to further validate the existence of the active species^32^. Detailed protocols for the probe degradation assays are provided in **Text S11**. Since the aforementioned degradation experiment was conducted under conditions distinct from those of the disinfection trials, terephthalic acid (TA) was subsequently introduced as an auxiliary probe to corroborate the formation of radical species under the same operational parameters used for viral inactivation^33^. The quantification of all probe compounds (NB, BA, DMOB, CBZ, TA) and the fluorescent TA-derived products was performed by high-performance liquid chromatography (HPLC, Agilent 1200, USA), with the detailed analytical parameters provided in **Table S4.**

## 3 RESULTS AND DISCUSSION

### 3.1 Effective Inactivation of Phi6 by EC disinfection

The complete inactivation (> 7-log) of Phi6 was achieved by EC disinfection when the cell voltage exceeded 4 V in 10 mM NaCl solution (**Figure 1a**). No bacteriophage plaque was observed in the effluent samples under these conditions. Interestingly, a sharp, ∼6-log shift in inactivation occurred as the cell voltage increased from 3 to 4V. This dramatic change might be associated with the generation of electrochemical reactive species, as discussed below.

**Figure 1.**
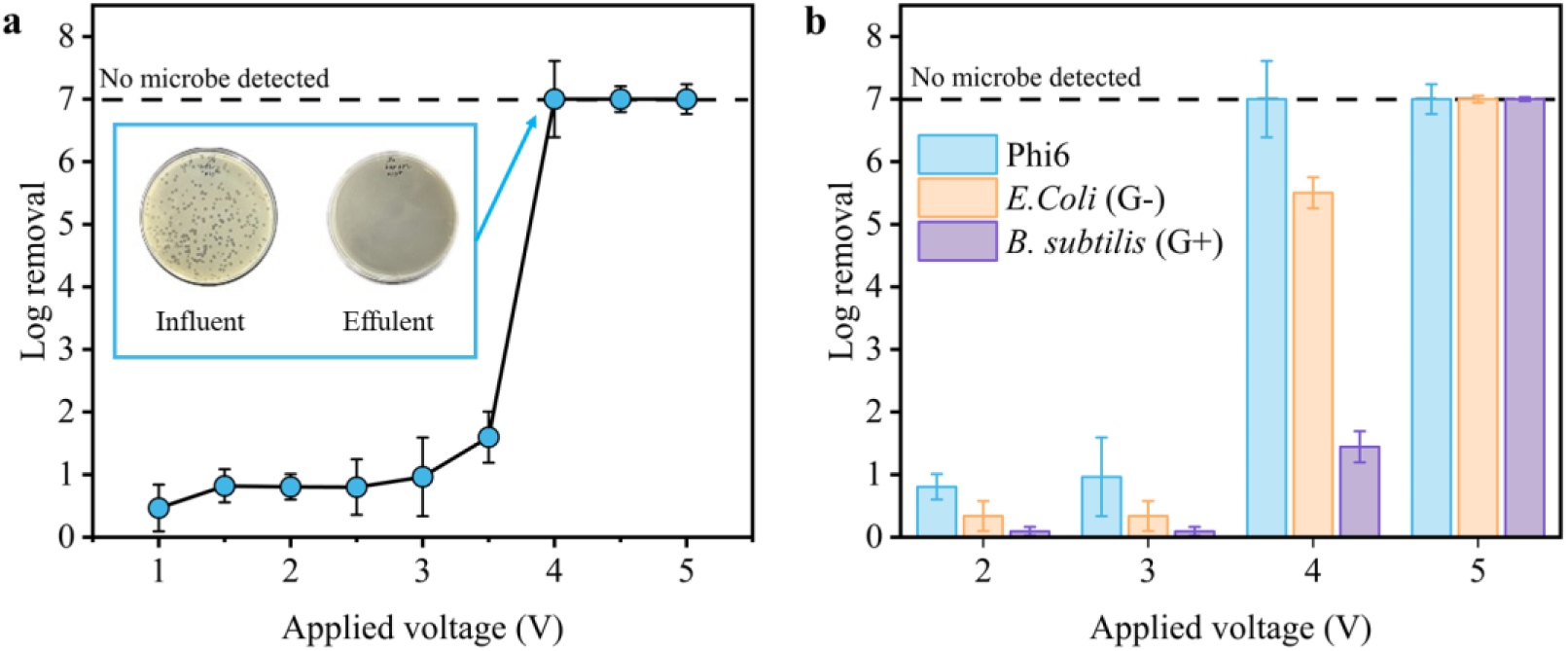
Inactivation efficacy of (a) Phi6 and (b) two typical bacteria, *E. coli* (G-) and *B. subtilis* (G+) by EC disinfection at different cell voltages.

A rank of relative resistance as Phi6 < *E. coli* < *B. subtilis* was represented under the EC disinfection (**Figure 1b**). At the cell voltage of 4 V, the loss of infectivity was 5.5-log for *E. coli* and 1.7-log for *B. subtilis*, both of which were lower than the 7-log reduction observed for Phi6. The high susceptibility of Phi6 may be attributed to the absence of protection from the cell wall and extracellular polymeric substances, as well as its limited self-repair capability when partially damaged^17^. Notably, this finding contrasts with prior studies reporting that non-enveloped viral surrogates (e.g., MS2, rAdeV) often exhibit greater tolerance to EC treatment than bacterial surrogates^25^. While in this study, as a representative of enveloped viruses, Phi6 shows significantly higher sensitivity than model bacteria. Indeed, this finding aligns with a substantial body of literature, which has collectively reported that enveloped viruses typically exhibit reduced tolerance to most disinfection or environmental processes, such as chlorination and solar irradiation^13^. Hence, as an effective supplement, our study confirms that enveloped viruses are also highly susceptible to EC disinfection, providing critical insights for evaluating EC as a broad-spectrum water disin-fection approach.

### 3.2 Severe damage to viral structures during EC disinfection

Convergent evidence from multiple molecular biological analyses collectively demonstrated the substantial damage to the structural proteins, envelope integrity, and RNA genome of Phi6 following EC disinfection. According to TEM observations, intact spherical structures of Phi6 were visible before disinfection (**Figure 2a, Figure S4**). However, after EC treatment, almost no intact Phi6 particles could be found across the entire TEM viewing field, with only some amorphous debris, suspected to be the remnants of disrupted viral particles. This is in stark contrast to previous studies, which have shown that discernible viral structures often persist even after levels of inactivation^14^. Hence, the complete disappearance of characteristic viral structures in our TEM observations suggests that EC disinfection caused thorough destruction of Phi6.

**Figure 2.**
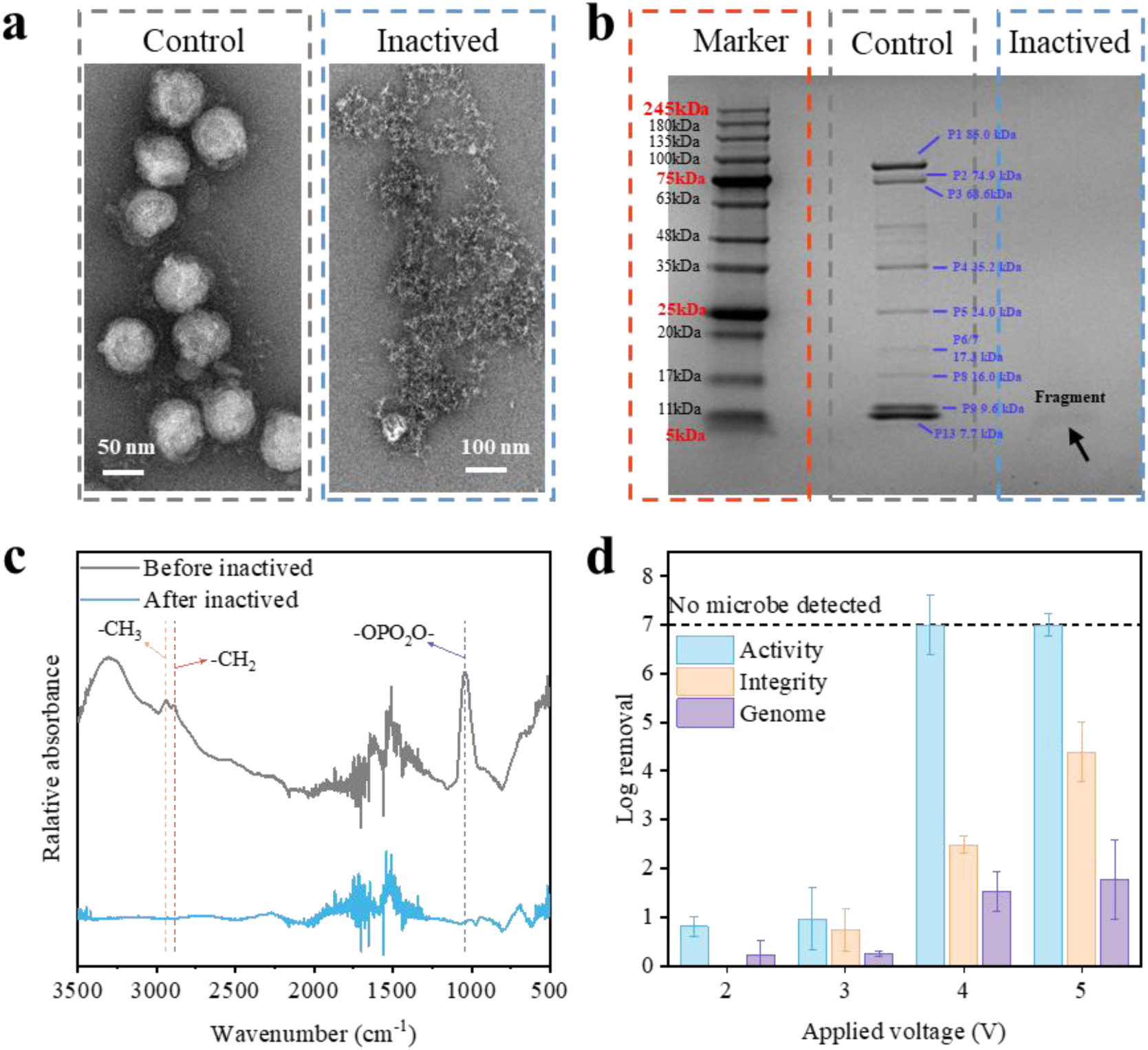
The damage to (a) the integrity of viral particles observed by TEM, (b) viral proteins by SDS-PAGE electrophoresis, and (c) lipid envelope membrane by FTIR of Phi6 under the EC disinfection. (d) The loss of infectivity, integrity, and RNA genome following EC disinfection under different applied cell voltages.

Concurrently, damage to the proteins of Phi6 was further assessed via SDS-PAGE electrophoresis. Before disinfection, a total of nine characteristic proteins belonging to Phi6 were detected, categorized as envelope proteins, capsid proteins, and functional proteins within the viral capsid (**Table S6**). After disinfection, a complete disappearance of the original nine viral protein bands was observed, leaving only very faint bands in the 5-10 kDa region (**Figure 2b**). This result reveals that nearly all Phi6 proteins were severely fragmented and degraded into small peptides (< 10 kDa) during EC disinfection. Moreover, it is noteworthy that proteins P2 and P7 are deeply buried within the viral particles, enclosed by both the envelope and capsid layers, affording them maximal protection from damage. The severe destruction of these deeply buried proteins indicates that EC disinfection either thoroughly disrupted the viral structure from the outside inward or generated reactive species capable of penetrating deeply to inflict damage. Furthermore, FT-IR spectroscopy was employed to evaluate the loss of lipids from the viral envelope before and after disinfection (**Figure 2c**). Before disinfection, the FT-IR spectrum showed apparent absorption peaks at 2941, 2887, and 1042 cm^-1^, corresponding to the -CH_3_, -CH_2_, and -OPO_2_O-functional groups of phospholipids, respectively. Additionally, multiple absorption peaks could be observed around 1250-2000 cm^-1^, corresponding to different secondary structures of proteins on the Phi6 envelope. After EC inactivation, the characteristic phospholipid peaks were almost completely absent, exhibiting severe damage to the Phi6 lipid envelope.

The loss of Phi6 infectivity, integrity (via PMA-qPCR), and RNA genome after EC disinfection was further compared as illustrated in **Figure 2d**. A clear order emerged when the applied voltage exceeded 3V (infectivity loss > integrity loss > genome damage), suggesting that damage to the genome during electrochemical disinfection requires the prior destruction of the viral capsid structure. With the cell voltage set as 4V and 5V, 99.690% (2.48-log) and 99.996% (4.38-log) of Phi6 lost their integrity, respectively. These results are consistent with the complete absence of intact viral particles observed by TEM (**Figure 2a; Figure S4**). Concurrently, the damage to Phi6 genome reached 1.5-log (∼97%) at both 4V and 5V. It is critical to emphasize that this genomic damage was quantified by direct RT-qPCR measurement without theoretical extrapolation (which would imply a >100-log reduction for the full genome)^29,34^. Furthermore, the RT-qPCR amplicon in this study was intentionally designed to be short (100 bp). Given that short amplicons are known to be detectable even in extensively degraded genomes^35,36^, the severe degradation observed here for this 100-bp target implies that the entire viral genome suffered extensive fragmentation. In sharp contrast, a previous study on Phi6 inactivation by FC and UV achieved ∼6-log infectivity reduction with only 10-50% damage to a 500-bp amplicon on the S segment—the same genomic region targeted in this study^29^. This striking disparity highlights that, despite achieving similar inactivation efficacy, EC disinfection inflicts markedly more severe genomic fragmentation.

In summary, the collective evidence from multiple analytical techniques leaves no doubt that electrochemical disinfection achieves thorough destruction of all viral compo-nents—from the outer envelope and capsid to the internal genome. Moreover, the degree of devastation observed here is rarely achieved by conventional disinfection methods.

### 3.3 Electrogenerated RCS as the Primary contributor of Virus Inactivation

Following the observation of extensive structural damage to Phi6, we attempted to identify the specific EC reactions responsible for viral inactivation (possible EC reactions illustrated in **Figure 3a**). The respective contributions of EC reactions and electric field effects were initially differentiated by applying frequency-specific alternating current (AC; 10³–10⁶ Hz), which has been proven to suppress electrolytic processes while maintaining the efficacy of electroporation sufficiently^37,38^. **Figure 3b** compares the inactivation efficacy under DC and AC (10^5^ Hz) treatments at equivalent voltages, revealing a distinct difference between the two application modes. Hence, these results demonstrate that EC reactions, rather than electric field effects, played the dominant role in the inactivation.

**Figure 3.**
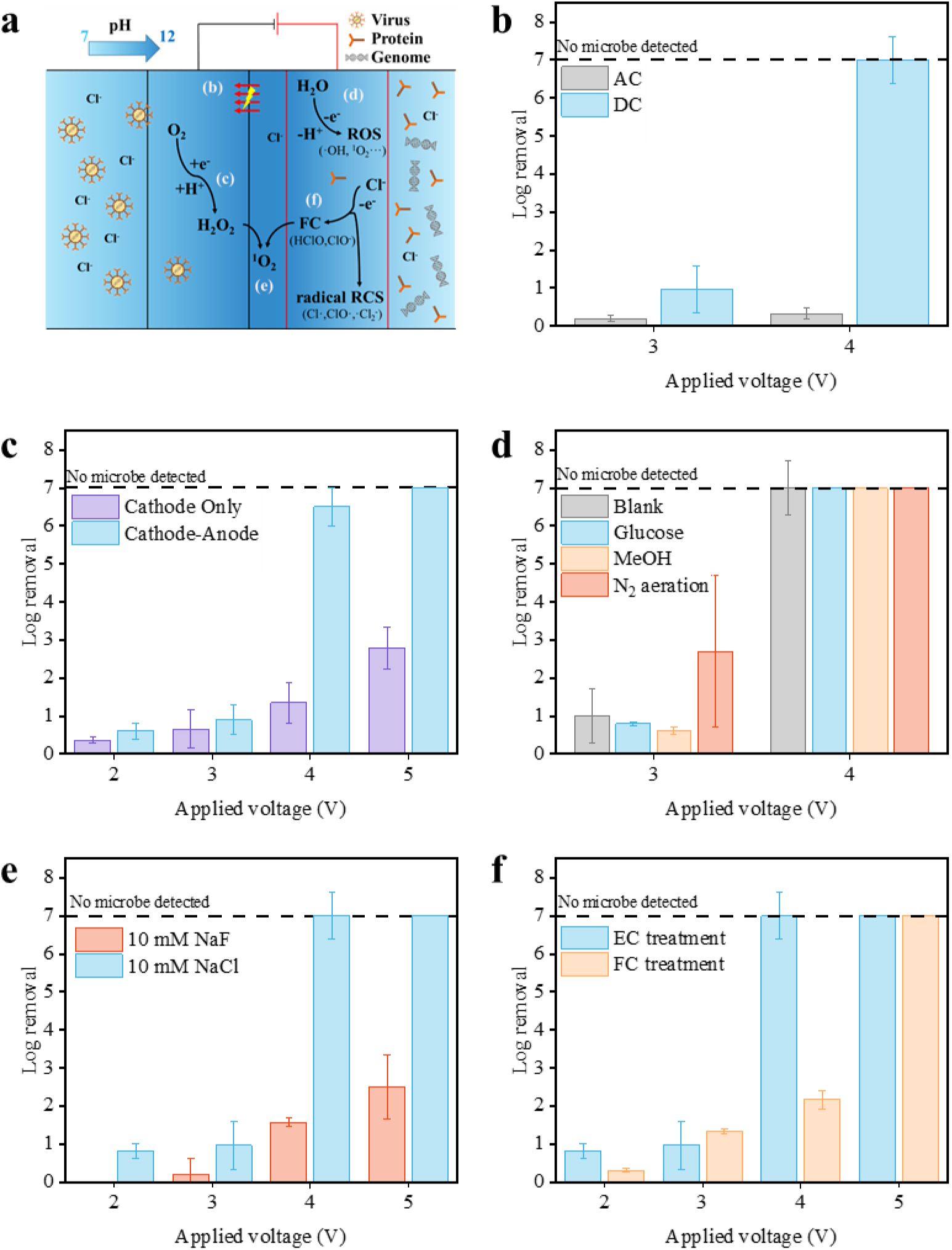
(a) Schematic diagram of main EC processes and the comparisons of Phi6 inactivation under standard condition by EC treatment with (b) high frequency AC for electric field effect, (c) cathode only for cathodic EC reactions, (d) MeOH masking and N_2_ aeration deoxidation for ·OH and ^1^O_2_, (e) 10 mM NaF solution for RCS generation, and (f) simulated FC solution for electrogenerated RCS.

To further elucidate the contributions of the cathodic versus anodic EC reactions, samples were collected from the intermediate chamber of the EC reactors (as illustrated in **Figure S2**) to assess the inactivation solely attributed to the cathode. As shown in **Figure 3c**, measurable inactivation occurred from the cathode treatment and increased with higher cell voltages. H_2_O_2_ was detected in the cathode effluent across voltage gradients, and the simulated tests (**Text S7**) suggested H_2_O_2_ partially accounted for the limited, cathode-driven Phi6 inactivation (∼1-log reduction, **Figure S5**). Additionally, the significant change of pH under high external voltage treatment (from 7 to 11, **Figure S6**) might also contribute to the additional inactivation of Phi6 (**Figure S7**). It is worth noting that the pH within the porous cathode will be even higher than that in the intermediate chamber. However, despite the partial inactivation achieved at the cathode, non-negligible gaps in the efficacy were still observed between the complete cathode-anode EC reactor and cathode-only treatment (**Figure 3c**). Therefore, it could be inferred that the anodic EC reactions may be more important for the Phi6 inactivation.

Based on the preceding analysis, the relative importance of different anodic EC reactions, including the anode-generated ROS and RCS, as well as the direct anode oxidation, was further quantified. Methanol (MeOH), a well-documented radical quencher, was employed to scavenge ·OH. Importantly, at the concentration (10 mM) used in this study, MeOH exhibited minimal toxicity to both Phi6 and its host bacteria. As shown in **Figure 3d**, the addition of MeOH did not significantly diminish the inactivation efficiency compared to the blank or the TOC-matched glucose control, suggesting that ·OH was not the major contributor to viral inactivation. Moreover, the role of another important ROS, ^1^O_2_, was further evaluated. Since most quenchers of ^1^O_2_ (e.g., NaN_3_) exhibit high biological toxicity, and the generation of ^1^O_2_ in EC systems primarily relies on the reduction of oxygen at the cathode (**Equations 2–5**), we instead suppressed the generation of ^1^O_2_ by removing dissolved oxygen (DO) via nitrogen (N_2_) purging (detailed protocols in **Text S9**). However, as shown in **Figure 3d**, DO removal did not lead to a reduction in the inactivation rate of Phi6—instead, it resulted in a slight increase. This indicates that ^1^O_2_ is also not a primary factor responsible for the inactivation of Phi6.

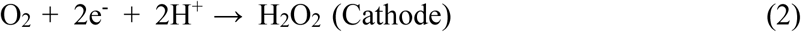

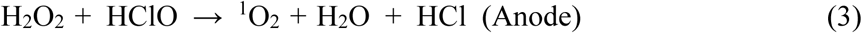

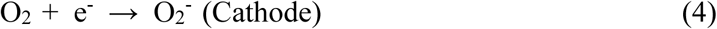

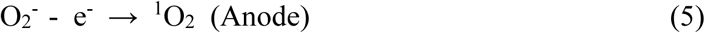

Furthermore, to investigate the contribution of RCS to the inactivation, NaCl in the influent feeding solution was replaced with NaF at an equivalent concentration to maintain similar conductivity. Following this change, a pronounced decline in the inactivation efficacy was observed (**Figure 3e)**, indicating that chlorides are essential for Phi6 inactivation. Moreover, when a pure 10 mM NaCl solution (without viruses) was supplied as the influent, free residual chlorine was detected in the EC reactor effluent, providing direct evidence of the generation of RCS via the EC reactions (**Figure S8**). As reported, enveloped viruses, such as SARS-CoV-2^39^ and Influenza^40^, as well as the model surrogates like Phi6, are highly susceptible to chlorine-containing disinfectants^22^. Typically, a relatively low con-centration-time product (CT value) of 0.03∼2.05 mg min L^-1^ was sufficient to achieve > 4-log of these enveloped viruses^22,39,40^. In this study, the estimated CT values of FC were 0.81 and 3.5 mg min L^-1^ at cell voltage of 4 V and 5 V (shown in **Figure S8**), which fall within this effective range. Consequently, these results indicate that the electrogenerated RCS played a crucial role in the inactivation of Phi6. Additionally, to further determine whether the direct anode oxidation also contributed to the inactivation, the individual contributions of anodic and cathodic EC reactions were re-evaluated under the 10 mM NaF electrolyte system. The results demonstrated a minimal contribution from direct anode oxidation (**Figure S9**). In summary, based on the comprehensive evidence presented above, it is reasonable to conclude that the inactivation of Phi6 during the EC disinfection was primarily driven by the generation of RCS through the anode EC reaction, with other EC reactions contributing minimally to the overall inactivation process.

Though Phi6 was inactivated by the RCS generated via EC reactions, it is important to note that the EC disinfection caused distinctly different damage to viruses compared to the conventional FC disinfection. As discussed in Section 3.2, and consistent with prior studies, damage to the viral genome by FC remains relatively limited^22,23,29^—even at high inactivation rates—and does not reach the severe level of disruption observed in this study. This difference suggests that the RCS generated during EC disinfections may possess higher reactivity than conventional FC treatment. More importantly, as shown in **Figure 3f**, the inactivation efficiency under 4V EC treatment exceeded that of simulated FC disinfection, despite identical total chlorine doses and CT values (for details, see **Text S10** and **Figure S8**). Collectively, these evidences indicate that RCS generated during EC disinfection is more reactive than sodium hypochlorite. Two theoretical mechanisms may explain these observations. On the one hand, given that the majority of EC processes involve interfacial reactions, microenvironments characterized by low pH values and high local RCS concentrations are likely to form near the anode surface. Within these microenvironments, the enhanced inactivation efficacy can be attributed to a higher effective RCS concentration and proportion of HClO compared to ClO⁻ at lower pH values, which has been extensively documented in prior studies^20^. On the other hand, it has been reported that the radical chlorine species, such as ·Cl, ·ClO, and ·Cl_2_-, can also be generated via the anode EC reactions^30,31^. These radical species often exhibit faster chemical reaction kinetics than conventional FC, particularly toward aromatic structures with electron-donating functional groups, which are commonly found in proteins and nucleic acids^41,42^. Therefore, the presence of ·Cl,·ClO, and ·Cl_2_- may also account for the substantial damage to viral proteins and genomes observed during the EC disinfection in this study. However, due to the considerable challenges associated with *in situ* characterization of interfacial reaction processes, the precise contributions of these two mechanisms to viral inactivation in EC systems require further investigation.

### 3.4 Verification of the presence of ·ClO, and ·Cl_2_-

Owing to the limitations imposed by biocompatibility, many validation experiments could not be conducted within the disinfection system. Therefore, we performed the ex-situ experiments using chemical probes to further investigate the presence of various radical species. As shown in **Figure 4a**, the degradation kinetics of NB, a well-established selective probe for ·OH, exhibited no discernible change upon the addition of 100 mM TBA. Similarly, the BA (a probe for ·OH and ·Cl) also showed negligible degradation under the EC treatment. The combined evidence indicates that the formation of ·OH and ·Cl was minimal under the EC treatment, which was further supported by the absence of typical spectral lines in the electron paramagnetic resonance (EPR) results (**Figure S10**). In contrast, both DMOB and CBZ underwent significant degradation during the EC treatment, and their observed kinetics (*k*_obs_) declined markedly with the presence of TBA. Given that these two probes could also react with ·ClO and ·Cl_2_-, these findings suggest the generation of ·ClO and ·Cl_2_- (**Equations 6-9**).

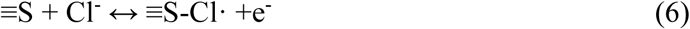

**Figure 4.**
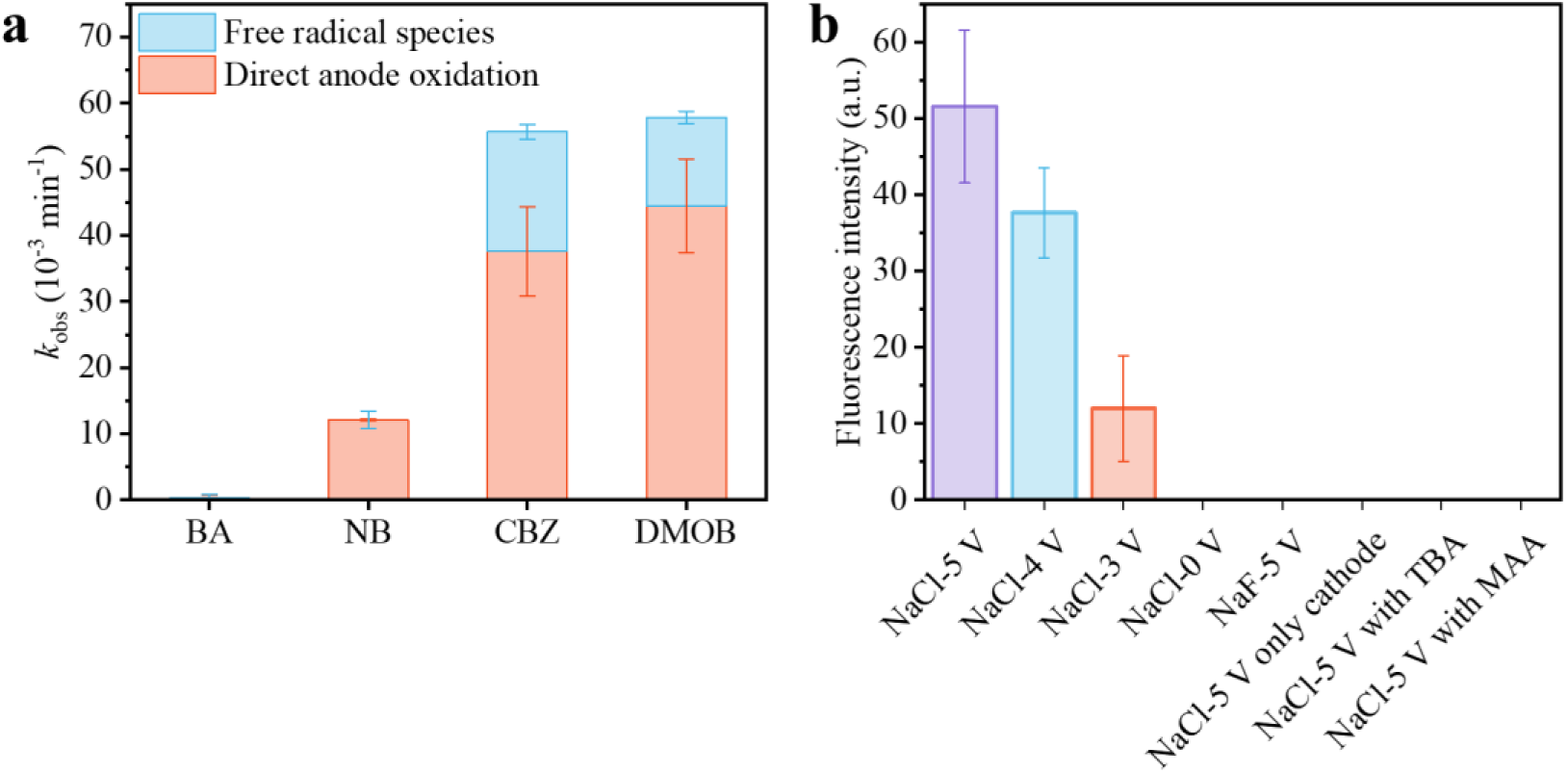
(a) The contribution of direct anode oxidation and free radical species to the removal kinetics of NB, BA, CBZ, and DMOB (b) the fluorescence intensity of TA transformation products under different conditions.

**Figure 5.**
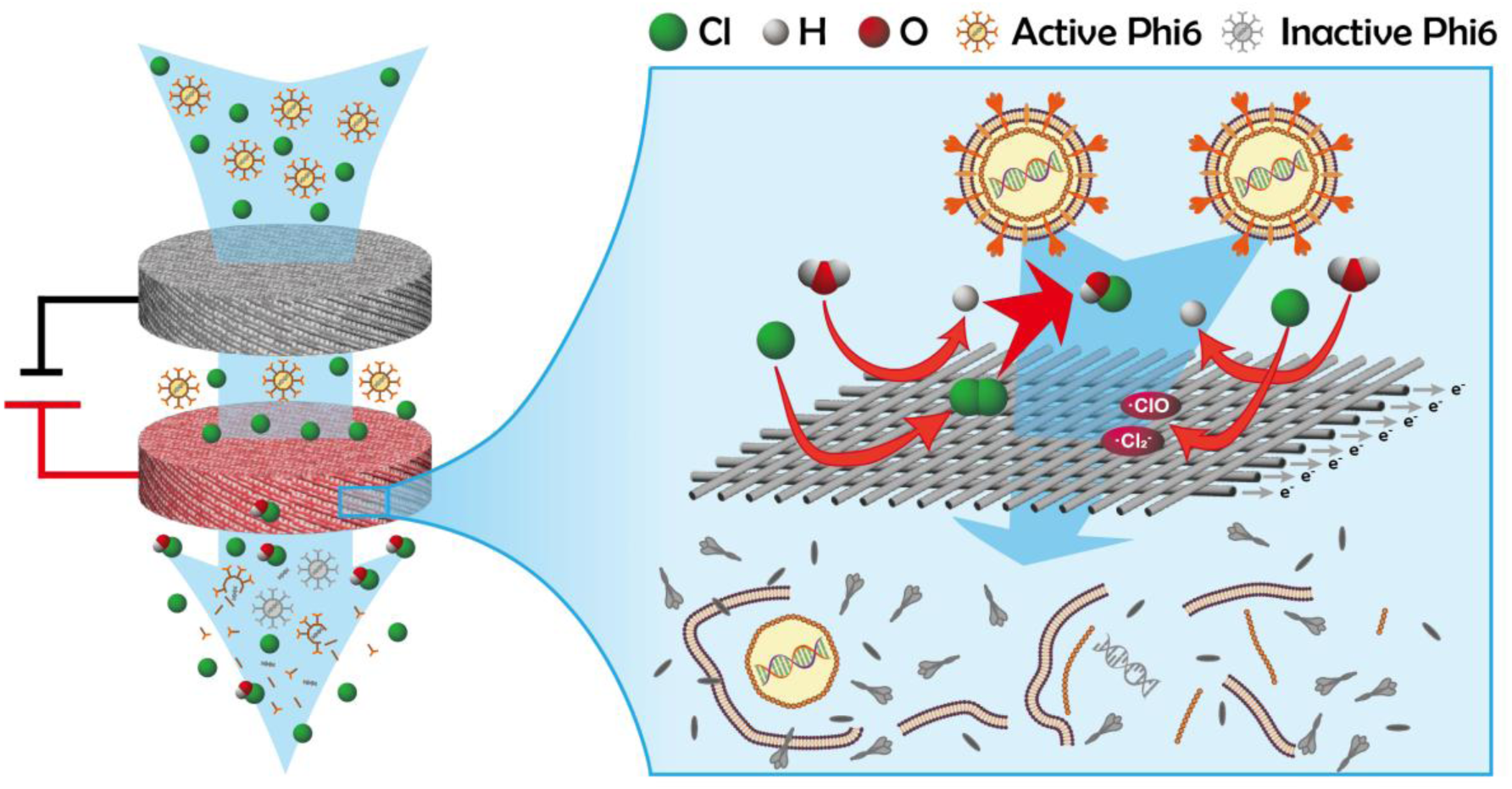
Schematic diagram illustrating the RCS electro-generation processes and damage mechanism for EC inactivation of enveloped viruses.

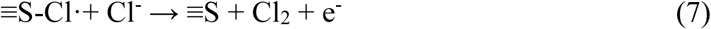

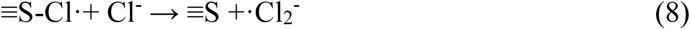

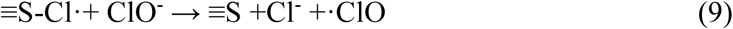

To further corroborate the formation of ·ClO and ·Cl_2_- during the EC disinfection, TA was employed as an additional probe and subjected to the same operating conditions as the disinfection assays. According to the results illustrated in **Figure 4b**, the complete absence of detectable fluorescence under the NaF electrolyte system further confirmed that ·OH was not generated. Interestingly, however, a pronounced fluorescence signal was observed in the effluent when NaCl was used as the electrolyte, and this fluorescence could be completely suppressed by radical scavengers (100 mM TBA and 10 mM MAA; **Figure 4b, Table S12**). Meanwhile, control experiments confirmed that neither FC, ^1^O_2_, nor the cathodically electrogenerated H_2_O_2_ and O_2_- could convert TA into fluorescent products (**Figure 4b** and **Text S13**). Although TA is usually regarded as a selective probe for the ·OH, an extensive body of experimental and theoretical studies has demonstrated that the ·ClO can likewise introduce a hydroxyl group to the benzene ring of benzoic acid derivatives via the radical adduct formation, yielding the fluorescent salicylic acid structure (as discussed in **Text S14**, **Figure S14**). To unambiguously attribute the observed fluorescence in the NaCl system to ·ClO rather than to the trace ·OH arising from the altered electrolyte environment (NaCl vs NaF), we simultaneously tracked the degradation of BA and CBZ under identical conditions. As shown in **Figure S15 and S16**, BA remained essentially inert, whereas CBZ exhibited significant degradation relative to the NaF control, corroborating the presence of ·ClO. Furthermore, assuming that all the fluorescent products formed via TA reacting with ·ClO exhibited the same quantum yield as 2-hydroxyterephthalic acid, the effective concentrations of electro-generated ·ClO were estimated to be 37.6 nM at 4V and 51.6 nM at 5V, respectively. This concentration level is approximately six orders of magnitude higher than that of the spiked Phi6 virus, sufficient to induce complete disruption of all the viral structures. Therefore, all the probe assays collectively confirm that the EC-generated ·ClO and ·Cl_2_- may lead to the thorough destruction of Phi6 during the dis-infection process.

## Supporting information

The MIQE checklist for this manuscript

The supplemental file containing the texts, figures and tables

## 4 ENVIRONMENTAL IMPLICATIONS

Waterborne viruses are increasingly recognized as an emergent threat to global water safety, and the EC disinfection stands out as a next-generation technology for addressing this challenge. This study bridges a critical knowledge gap by demonstrating that EC treatment achieves both rapid inactivation and complete viral structure disintegration. These findings provide crucial evidence for assessing the reliability of EC disinfection and support its broader implementation.

Meanwhile, our results elucidate the decisive role of electrogenerated RCS in viral inactivation. Given that chloride is the predominant and most abundant anion in most natural waters and wastewater, this mechanism underscores the high feasibility and promising application prospects of EC disinfection for real-world water disinfection. However, the generation of RCS also indicates that the EC disinfection is not an “absolutely green” process. Consequently, future studies should take efforts to investigate the formation profiles of disinfection by-products and the secondary risks arising from the EC disinfection in real complex water matrices.

Furthermore, while quantifying the precise contribution of radical RCS was challenging due to the complexity of the EC system and biocompatibility constraints, our findings nonetheless demonstrate that they may accelerate the viral inactivation kinetics and exacerbate viral structural disruption. Notably, radical RCS are not unique to the EC system, but are also generated in other water treatment processes, such as the UV/Chlorine disinfection. Therefore, a systematic understanding and targeted utilization of radical RCS in such processes could substantially strengthen the control of waterborne pathogens and associated biological threats, particularly during public health emergencies.

## 5 ASSOCIATED CONTENT

Schematic of Phi6 and EC reactor; additional experimental protocol details and results of microbes culturation, control tests, and probes tests; characteristics of CFF electrode; sequences of primers and probes of Phi6; basic information of Phi6 protein profiles; summary of previous electrochemical inactivation of viruses; HPLC conditions; pseudo-first-order reaction kinetics fitting of probes.

## 6 ACKNOWLEDGMENTS

The authors gratefully acknowledge the financial support from the Major Program of the National Natural Science Foundation of China (No. 52091543).

