## Supplementary material for "Rapid and Complete Inactivation of Enveloped Viruses by Electrochemical Disinfection: Unraveling the Contribution of Reactive Chlorine Species": The supplemental file containing the texts, figures and tables

**This supporting information has 49 pages, containing 15 associated texts, 22 figures and 12 tables.**

**Figure S16** The probe tests of CBZ in the flow-through EC reactor in 10 mM NaCl and NaF. . 31

**Text S1. Materials and chemicals.**

Ethanol, NaCl, NaF, $Na_2SO_4$, $KH_2PO_4$, D-(+)-Glucose, $NH_4Cl$, nitrobenzene (NB), benzoic acid (BA), 2,4-dimethoxybenzene (DMOB), and carbamazepine (CBZ), 5,5-dimethyl-1-pyrroline N-oxide (DMPO), disodium terephthalate (2Na-TA), 2-hydroxyterephthalic acid (hTA), 2-methallyl alcohol (MAA), methanol (LC-MS), and $Na_2HPO_4$ were purchased from Aladdin Biochemical Technology Co., Ltd., China. NaClO standard solution (30 g $L^{-1}$), $Na_2S_2O_3 \cdot 5H_2O$, $Na_2HPO_4 \cdot 12H_2O$, $MgSO_4 \cdot 7H_2O$, and tert-Butanol (TBA) were purchased from Shanghai Macklin Biochemical Technology Co., Ltd., China. Tryptone soy broth (TSB), agar powder, and 1×PBS (pH 7.2-7.4, 0.01 M) were purchased from Beijing Solarbio Science & Technology Co., Ltd., China. Eosin-methylene blue agar (EMB) and 30% $H_2O_2$ solution were purchased from Shanghai Bio-way Technology Co., Ltd., China. Acetone was purchased from Sinopharm Chemical Reagent Co., Ltd., China. Carbon fiber felt (CFF) was purchased from Liaoning Jingu Carbon Materials Co., Ltd, China, and its physical and chemical properties are shown in **Table S1**.

All solutions were prepared using ultra-pure water (resistivity ≥ 18.2 MΩ·cm) produced by a Dura water purification system. Tryptic Soy Broth (TSB) liquid medium, TSB semi-solid medium, and Eosin Methylene Blue (EMB) agar medium were prepared according to the manufacturer's instructions. M9 culture medium is prepared by dissolving 1 g $KH_2PO_4$, 0.3 g $Na_2HPO_4$, 4 g D-(+)-Glucose, 0.5 g $NH_4Cl$, and 1 mL 1 M $MgSO_4$ into 1 L ultra-pure water, followed by sterilization through a 0.45 μm membrane filter.

Phi6 and its host *Pseudomonas syringae* (*P. syringae*) (ATCC 21781) were purchased from American Type Culture Collection (ATCC). *Escherichia coli* (*E.coli*, G-) (ATCC 15597) and *Bacillus subtilis* (*B. Subtilis*, G+) (CMCC(B)63501) were purchased from the Shanghai Microbiological Culture Collection Co., Ltd, China.

**Text S2. The culturation, purification, and quantification of *E.coli* and *B.subtilis*.**

*E. coli* ($G^{-}$) and *B. subtilis* ($G^{+}$) were selected as representative Gram-negative and Gram-positive bacterial surrogates, respectively, for resistance comparison with Phi6. Briefly, each bacterium was propagated in TSB broth at 37 °C for 24 h. The cultures were then centrifuged at 8000×g for 15 min, and the resulting pellet was resuspended in 10 mM NaCl solution. This centrifugation–resuspension cycle was repeated three times to thoroughly remove residual organic matter from the culture medium, yielding a bacterial stock solution with a concentration of approximately $10^9$ CFU $mL^{-1}$. The stock was diluted 100-fold with 10 mM NaCl before inactivation assays, which were conducted under the same conditions as those used for Phi6 (Details in **section 2.2**). The concentrations of *E. coli* and *B. subtilis* before and after disinfection were determined using the dilution plating method on EMB agar and TSB agar, respectively, and were quantified using the same formula as applied in the Phi6 inactivation tests.

**Text S3. Protocol for RT-qPCR assay and quantification.**

The primer/probe set (listed in Table S2) used for Phi6 genome quantification (listed in Table S2) was previously described. The specificity of this set was assessed by agarose gel electrophoresis, with the Phi6 genome serving as a positive control and the host's total RNA/DNA as a negative control (Data not provided).

For the absolute quantification, recombinant PUC57 plasmid DNA containing the amplicon sequence was synthesized by Sangon. The lyophilized plasmid was resuspended in TE buffer (10 mM Tris, 1 mM EDTA, pH = 8.0) and its mass concentration was determined using the Qubit HS dsDNA Assay (Thermo Fisher, Q32854). Subsequently, it was linearized with the SspI restriction enzyme (Sangon, B600763), diluted 10^6-fold, and quantified absolutely on a digital PCR platform (dPCR) using the same primer-probe set as for RT-qPCR. The dPCR quantification was conducted on the DropXpertS6 (BIORAIN) platform by Sangon. The quantification result was deemed acceptable only if the dPCR-measured concentration was within 20% of the theoretical value calculated from the mass concentration. It should be noted that, since Phi6 is a double-stranded RNA virus, using a plasmid standard for absolute quantification may introduce systematic errors. However, given the practical difficulties in synthesizing high-purity dsRNA, the plasmid was still selected as the standard for quantification. The qualified plasmid was diluted to $10^7$ copies/μL, aliquoted, and stored at -80°C. Before use, an aliquot was thawed and serially diluted in TE buffer (1-$10^5$ copies/μL) to prevent the impact of repeated freeze-thaw cycles on the quantified value.

To ensure efficient reverse transcription of the dsRNA genome of Phi6 during the one-step RT-qPCR, the nucleic acid samples were firstly heat-denatured at 65°C for 5 min in a PCR instrument, immediately chilled on ice, and all subsequent PCR preparation steps were carried out on ice. The

RT-qPCR was performed in a 20 μL reaction volume, containing 10 μL of 2× FastKing One Step Probe RT-qPCR Mix, 0.8 μL of 25× FastKing Enzyme Mix, 1 μL each of the 10 μM forward and reverse primers, 0.5 μL of a 10 μM probe, 5 μL of template, and 1.7 μL of RNase-free water. For each sample, three parallel tests were set to reduce the quantification bias. For each plate, a negative control and a dilution series of quantification standards (from 1 to $10^5$ copies/μL) were included. The RT-qPCR thermocycling protocol was as follows: reverse transcription at 50°C for 30 min; initial denaturation at 95°C for 5 min; then 40 cycles of denaturation at 95°C for 15 sec and annealing/extension at 60°C for 30 sec. All the RT-qPCR reactions were performed on the QuantStudio 5 qPCR platform (Thermo Fisher).

For quality control, every RT-qPCR assay was validated according to three criteria: 1) the standards displayed a sigmoidal amplification curve with no amplification in the negative control; 2) the standard curve exhibited an $R^2 > 0.99$ with an efficiency of 90–110%; 3) amplification of the 1 copy/μL standard was detected in at least one of the three technical replicates. Upon passing the quality control criteria, the CT values from each reaction were automatically calculated by the Quanstudio software. The mean CT value from the three replicates for each sample was employed to determine its concentration via the standard curve.

**Text S4. Protocol for propidium monoazide (PMA) staining and photo-inactivation.**

For PMA staining, 0.5 µL of PMA (20 mM; Biotium, 40019) was added to 1 mL of effluent sample from the EC reactor to achieve a final concentration of 100 µM. The mixture was vortexed briefly and incubated in the dark at room temperature for 5 minutes. After staining, residual PMA was photo-inactivated by exposure to a 500 W halogen tungsten lamp (Philips, QVF135). During irradiation, the lamp was positioned 20 cm above the samples, which were kept on ice to minimize thermal degradation. Following PMA inactivation, 200 µL of each sample was subjected to nucleic acid extraction.

Given that the optimal concentration of PMA for distinguishing intact from damaged viral particles may vary among species, we assessed its applicability to Phi6 using $10^7$ PFU of freshly cultured Phi6 as a negative control and $10^7$ copies of extracted Phi6 nucleic acids as a positive control. The validation results, summarized in **Table S3**, show that PMA treatment had no effect on fresh Phi6 culture but completely suppressed amplification of free Phi6 nucleic acids. These findings confirm that PMA treatment followed by photo-inactivation under the specified conditions effectively differentiates between intact and damaged Phi6 viral particles.

**Text S5. Optimized protocol for preparing high-purity viral stocks.**

To prepare highly purified Phi6 stocks critical for the analysis of SDS-PAGE electrophoresis, TEM, and FT-IR, we optimized both the cultivation and purification methodologies. Briefly, Phi6 was propagated using M9 medium instead of TSB. As M9 is a protein-free medium, this modification will significantly reduce the amount of external protein. The procedures for propagating and harvesting *P. syringae* and Phi6 in M9 medium were identical to those used in TSB medium (as described in Section 2.1). After the concentration by ultrafiltration, the Phi6 stock was further subjected to ultracentrifugation to remove the residual cell debris. Briefly, a 10-40% (w/v) step sucrose gradient (8 mL for each step) was prepared in a 38.5 mL open-top polypropylene tube (Beckman Coulter, 369650). The ultrafiltration-concentrated Phi6 stock was carefully layered onto the top of the 10% (w/v) sucrose gradient, followed by the addition of PBS to fill the remaining volume of the tube. The loaded tube was then placed in an SW 32 Ti swinging-bucket rotor (Beckman Coulter, 369650) and centrifuged at 65,700 × g for 1.5 h using an Optima XPN-100 ultracentrifuge (Beckman Coulter). Following ultracentrifugation, the opalescent band at the 30–40% sucrose interface was carefully aspirated with a long-needle syringe and diluted in 100 mL of PBS. To remove residual sucrose, the sample was subjected to three additional rounds of ultrafiltration using a new 30-kDa molecular weight cutoff centrifugal device (UFC703008, Millipore). The purity of the final Phi6 preparation was assessed by SDS-PAGE, which revealed a near-exclusive presence of Phi6-specific protein bands. The concentration of the purified Phi6 stock was measured using the double-layer agar plaque assay. The freshly quantified stock was utilized directly in EC disinfection experiments to avoid any freeze-thaw cycles.

**Text S6. Protocols for Sodium Dodecyl Sulfate-Polyacrylamide Gel (SDS-PAGE) Electrophoresis, Transmission Electron Microscopy (TEM), and Fourier Transform Infrared Spectroscopy (FT-IR) Assays.**

For the SDS-PAGE electrophoresis, 40 μL of each sample, collected both before and after disinfection (with post-disinfection samples being concentrated as described in Section 2.4), were mixed with 10 μL of 5×SDS-PAGE loading buffer (Solarbio, P1040). The mixture was heated in a boiling water bath for 10 minutes, followed by centrifugation at 12,000 × g for 2 minutes. The entire supernatant was then loaded into a single well of a precast 8–20% SDS-PAGE gel (Solarbio, PG82010). Given the large loading volume, the samples were separated by an empty lane to prevent potential cross-contamination. Electrophoresis was carried out at a voltage of 120 V using a Mini-PROTEAN Tetra Vertical Electrophoresis Cell (Bio-Rad). Following the electrophoresis, the PAGE gel was first washed three times using DI water, and then stained with the Coomassie Blue solution (Sangon, E607056) with microwave-assisted heating. Subsequently, the background stain was removed by destaining with a solution containing acetic acid and methanol, also under microwave heating, until the protein bands became clearly visible. The stained gel was visualized and photographed using a Gel Imaging System (Tanon 5200).

For the TEM observation, 4 μL of each sample was applied onto a glow-discharged, ultrathin carbon-film copper grid and allowed to adhere at room temperature for 1 minute. The excess liquid was then blotted away using filter paper. Subsequently, the grid was inverted (with the virus-loaded side facing down) and sequentially rinsed by gently blotting it on three separate 40-μL droplets of 2% uranyl acetate solution, for 1 minute on each droplet. Following this, the grid was placed with the sample side facing up and air-dried for 1 minute. Any residual uranyl acetate solution was carefully removed by blotting with filter paper, and the grid was then stored in a grid box to airdry completely at ambient temperature. The prepared samples were finally imaged using a TEM (JEM-2100Plus, JEOL Ltd., Japan) operating at an accelerating voltage of 200 kV, with magnifications ranging from 4,000× to 200,000×.

For the FT-IR analysis, a 20 μL aliquot was frozen at -80°C and lyophilized for 24 h. The resulting lyophilized material was resuspended in 10 μL of deuterium oxide ($D_2O$) and directly applied onto an ATR crystal, where it was nitrogen-dried again. FT-IR spectroscopy was performed using an infrared spectrometer (Nicolet™ iS50 FTIR Spectrometer, Thermo Fisher) over a wavenumber range of 500–4000 $cm^{-1}$.

**Text S7. The inactivation tests of the equivalent $H_2O_2$ solution.**

To determine the contribution of cathode-generated $H_2O_2$ to virus inactivation, an equivalent $H_2O_2$ solution was prepared with the same $H_2O_2$ concentration and pH of the cathodic effluent. Phi6 stock was then spiked into the solution for an initial concentration of $10^7$ PFU $mL^{-1}$. 1 mL samples were taken out and immediately quenched by 1 mL 0.1 M $Na_2S_2O_3$ solution at 1, 2, 3, 4, and 5 min, respectively. The residual infectious titer was subsequently quantified using the double-layer agar plaque assay.

**Text S8. The inactivation tests of Phi6 at different pH values.**

To evaluate the contribution of pH changes to virus inactivation, the log-reduction of Phi6 was measured across a range of pH conditions (from 2 to 12). Solutions of 10 mM NaCl were adjusted to target pH values using 1 M HCl or NaOH. Phi6 stock was spiked into each solution to achieve an initial concentration of $10^7$ PFU $mL^{-1}$. After a 5-minute incubation at room temperature, an equal volume of 10× PBS buffer was added to neutralize the pH to 6–8. The residual infectious titer was subsequently quantified using the double-layer agar plaque assay.

**Text S9. Dissolved Oxygen Removal for Singlet Oxygen($^1O_2$) Suppression**

To suppress the generation of $^1O_2$, dissolved oxygen was removed via nitrogen ($N_2$) aeration. Briefly, 500 mL of 10 mM NaCl solution was placed in a 500 mL serum bottle and aerated with nitrogen below the liquid surface for 30 min. Subsequently, the headspace was purged for an additional 10 min to eliminate residual oxygen. The bottle was immediately sealed after sparging to prevent reoxygenation. Before disinfection, a Phi6 stock solution was spiked into the deoxygenated NaCl solution via syringe injection. The mixture was thoroughly vortexed, and then the bottle was uncapped. Disinfection assays were completed within 30 minutes to minimize reoxygenation. Following disinfection, the dissolved oxygen (DO) concentration was measured and found to be below 0.3 mg $L^{-1}$, confirming that the experiment was unaffected by reoxygenation.

**Text S10. Evaluation of Phi6 inactivation by free chlorine (FC) in a simulated anodic solution.**

The simulated solutions were prepared by diluting the NaClO stock (~5000 mg Cl $L^{-1}$) with 10 mM NaCl solution to different FC concentrations, matching the FC generation capacity of the EC reactor when fed with 10 mM NaCl solution in the absence of viruses (Figure S8). The inactivation assays were performed by mixing the simulated solution with Phi6 stock to achieve a final titer of $10^7$ PFU $mL^{-1}$, followed by continuous stirring for 2 minutes to simulate the hydraulic retention time (HRT) from the porous anode to the outlet pipe end of the EC reactor. The reaction was then quenched with 1 mL of 0.1 M $Na_2S_2O_3$ solution, and the residual concentration of infectious viruses was determined through double-layer agar plaque assay.

**Text S11. Multi-probe degradation assay for the detection of hydroxyl radical (·OH), chlorine radical (·Cl), dichlorine radical (·Cl), and chlorine oxide radical (·ClO).**

A set of probe compounds—NB, BA, DMOB, and CBZ was employed to identify the generation of ·OH, ·Cl, $\cdot Cl_2^-$, and ·ClO during the EC treatment[1,2]. Due to the high volatility of NB and DMOB, the probe experiments were conducted in an electrochemical cell equipped with a three-electrode system. CFF and Ti sheet with a diameter of 40 mm served as the working electrode and counter electrode, respectively, and Ag/AgCl electrode acted as the reference electrode. The distance between the reference electrode and the working electrode was approximately 2–3 mm. The electrolyte consisted of 200 mL of 10 mM NaCl solution. The anode potential was controlled at 2.5 V vs Ag/AgCl using an electrochemical workstation (PGSTAT302N, Autolab, Switzerland), matching the potential of the CFF anode in the EC reactor (**Figure S17**). To distinguish the contribution of anodic direct oxidation (ADO) to probe removal, 100 mM tert-butyl alcohol (TBA) was additionally introduced as a scavenger for free radicals (**Equation S1-2**).

$$\boldsymbol{k}_{\mathbf{ADO}} = \boldsymbol{k}_{\mathbf{obs,with\ TBA}} \tag{S1}$$

$$\boldsymbol{k}_{\mathbf{free\ RSs}} = \boldsymbol{k}_{\mathbf{obs,without\ TBA}} - \boldsymbol{k}_{\mathbf{obs,with\ TBA}} \tag{S2}$$

Based on the second-order reaction rate constants of CBZ and DMOB with ·ClO and ·Cl2⁻ (**Table S7**), the steady-state concentrations of ·ClO and $\cdot Cl_2^-$ were estimated by the non-negative least squares method, and were $6.32\times10^{-14}$ M and $3.41\times10^{-14}$ M, respectively. In addition, a control group with 5.25 mM $Na_2SO_4$ (with conductivity matched to 10 mM NaCl) was employed to assess the contribution of chloride. As summarized in **Table S8** and **Figure S22**, the degradation rate constants of the four probes in the $Na_2SO_4$ solution were comparable to those observed in the NaCl solution with TBA, indicating the absence of ·OH generation at the CFF anode.

**Text S12. The electron paramagnetic resonance (EPR) Assay.**

DMPO was employed as a spin trap agent to capture ·OH during ESR tests.[1,3,4] The EPR measurements (JEOL-FA200, Japan) were conducted in batch mode using a 50 mL EC cell with stirring at 300 rpm. Before each test, 40 mL of 100 mM NaCl solution was electrolyzed at 5 V for 2 minutes to eliminate residual contaminants until no background signal was detected. Subsequently, DMPO was introduced at a final concentration of 25 mM.[1] After electrolyzing for 1 minute at the target voltage, samples were immediately collected for EPR analysis. The instrument parameters were set as follows: center field 3360 G, sweep width 150 G, microwave frequency 9.43 GHz, microwave power 1 mW, modulation frequency 100 kHz, sweep time 120 s, and time constant 0.1 s. To enable consistent comparison of signal intensities across different conditions, the manganese oxide reference peaks were retained in all spectra.

**Text S13. The probe assay of terephthalic acid (TA) in the flow-through EC reactor.**

TA is one of the most recommended ·OH probes in electrochemical systems.[5,6] TA was often used as a quantitative or semi-quantitative probe for ·OH in chlorine-free systems ($Na_2SO_4$[7], $NaClO_4$[3,8,9] and $KH_2PO_4$[5,10]) for the selective reaction with ·OH to form fluorescent product 2-hTA[5,6]. Due to the advantages of non-volatility and high detection sensitivity of its fluorescent product, 100 μM TA was added in the influent to in situ identify the possible free radical species in the EC reactor.

However, the solubility of TA in water is strongly dependent on the solution pH, with poor solubility under acidic conditions. In the EC reactor, as shown in **Figure S6**, the pH varies drastically along the flow channel. Therefore, disodium terephthalate (2Na-TA) with higher solubility was used as a substitute for TA, and its selectivity for •OH was verified in a traditional Fenton system. As shown in **Figure S12,** the ·OH generation from the Fenton reaction was induced by the addition of $Fe^{2+}$, resulting in the fluorescence intensity (FI) significantly increasing. Moreover, the HPLC retention time of FI from 2Na-TA was the same as that of hTA, indicating the rationality of using 2Na-TA as a substitute for TA (**Figure S11**). Hence, 2Na-TA and TA were not distinguished in the main text and subsequent paragraphs in the support information. In addition, as shown in **Table S9**, TA can also be converted into fluorescent products in pure $H_2O_2$ solution, which may be due to the generation of a small amount of ·OH during the self-quenching process of $H_2O_2$[11], further verifying the good response ability of TA to ·OH.

The selectivity of the fluorescence signal for HClO and $^1O_2$ was also verified. As shown in **Table S10 and Table S11**, no fluorescence signal was observed under conditions of varying ionic strengths, pH values, and free chlorine (FC) concentrations, indicating that free chlorine cannot react with TA to form fluorescent products. The standard definition method was adopted to generate $^1O_2$ by adding NaClO to the $H_2O_2$ solution[12,13], and previous studies have shown that the conversion efficiency of this reaction is close to 100%.[12] However, the fluorescence signal of TA weakened with the increase in the amount of NaClO added (**Figure S13**), i.e., the TA fluorescence signal was negatively correlated with the $^1O_2$ concentration. The fluorescent products should be derived from residual $H_2O_2$, which indirectly proves that $^1O_2$ cannot induce the formation of fluorescent products such as hTA.

**Text S14. The inference of products from the reaction between TA and ·ClO.**

According to previous studies, fluorescent signals from TA oxidation products have been observed even in 100 mM NaCl solutions, despite the general understanding that such high concentrations of $Cl^-$ and electrogenerated $ClO^-$ can almost completely quench ·OH at the electrode surface and convert it into ·ClO[14]. Moreover, based on the current theoretical calculations, ·ClO primarily undergoes radical adduct formation with aromatic compounds, especially carbonylated benzene derivatives [15,16]. For -COOH/-$COO^-$ substituted benzenes, the reaction occurs mainly at the ortho and para-positions. In the case of terephthalic acid, adduction can only take place at the ortho-position. According to existing computational studies, the spin density on the oxygen atom of ·ClO is 0.78887[17], which is higher than that on the chlorine atom (0.2113[17]， 0.301[18]), indicating that the unpaired electron is predominantly localized on oxygen. Consequently, the oxygen atom is the primary site involved in radical adduction and bond formation[15,19]. Moreover, due to the lower bond dissociation energy of Cl-O (203 kJ/mol) compared to that of O–H (467 kJ/mol)[20]， the adduct may undergo subsequent dechlorination or radical rearrangement, ultimately leading to the formation of benzoquinone, phenol, or chlorophenol derivatives[15,16]. Therefore, as illustrated in **Figure S14**, it is theoretically plausible that during the reaction between ·ClO and TA, intermediates such as 2-chloro-3-hydroxyterephthalic acid or 2-hydroxyterephthalic acid (fluorescent) could be generated via the pathways mentioned above. Based on theoretical calculations using the recently developed FLuorophore design Acceleration ModulE (FLAME), it has been proposed that 2-chloro-3-hydroxyterephthalic acid may also exhibit fluorescence[21].

**Text S15. The electrochemical characterization of carbon fiber felt (CFF) electrodes.**

Linear sweep voltammetry (LSV) and cyclic voltammetry (CV) experiments were performed with a three-electrode system in 40 mL solution by Autolab PGSTAT302N (Metrohm, Switzerland). CFF and Ti sheet with the same area (20 mm×20 mm) served as the working electrode and counter electrode, respectively, and Ag/AgCl electrode acted as the reference electrode.

LSV was employed to characterize the anode direct oxidation of masking agents on the CFF anode. The experimental conditions were set as follows: scan rate of 5 $mV\cdot s^{-1}$, stirring rate of 300 rpm, and electrolyte solution of 10 mM NaCl (consistent with the EC reactor system). As shown in **Figure S19**, tert-butanol (TBA) did not undergo direct oxidation on the CFF electrode surface, while methacrylic acid (MAA) could bind to the active species on the electrode surface and be directly oxidized (**Figure S20**).

CV was used to investigate the effects of different probes and quenchers on the chlorine evolution reaction (CER) at the CFF anode, with a scan rate of 50 $mV\cdot s^{-1}$. As shown in **Figure S21**, when 100 mM NaCl was used as the electrolyte, a reduction peak appeared around 0.5 V vs Ag/AgCl, and the peak current increased with increasing scan number. In contrast, in a 50 mM $Na_2SO_4$ solution with similar conductivity, this reduction peak disappeared, and the anodic current decreased significantly. Additionally, introducing stirring during CV measurements could notably suppress the reduction peak. Based on these results, it is inferred that the reduction peak is related to the electrogenerated free chlorine accumulated inside the porous CFF electrode during CV tests. Subsequently, two masking agents, 100 mM TBA and 10 mM MAA, were added to the electrolyte solution, as shown in **Figure S22**. TBA had no inhibitory effect on chlorine evolution, whereas MAA significantly inhibited the reaction. This suggests that TBA can only react with free radical species, while MAA can also mask the adsorbed chlorine radicals ($\cdot Cl_{ads}$) on the electrode surface.

**Figure**

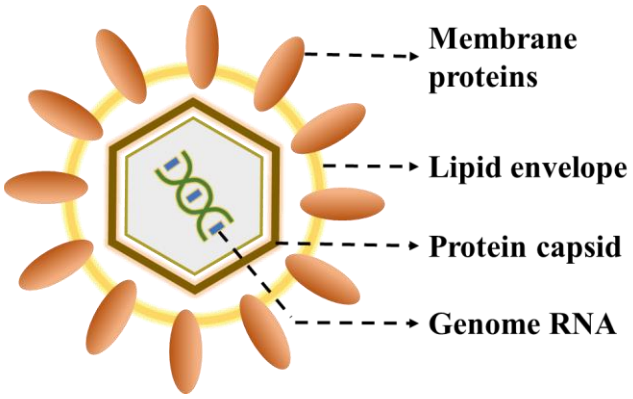

**Figure S1** The schematic diagram of Phi6.

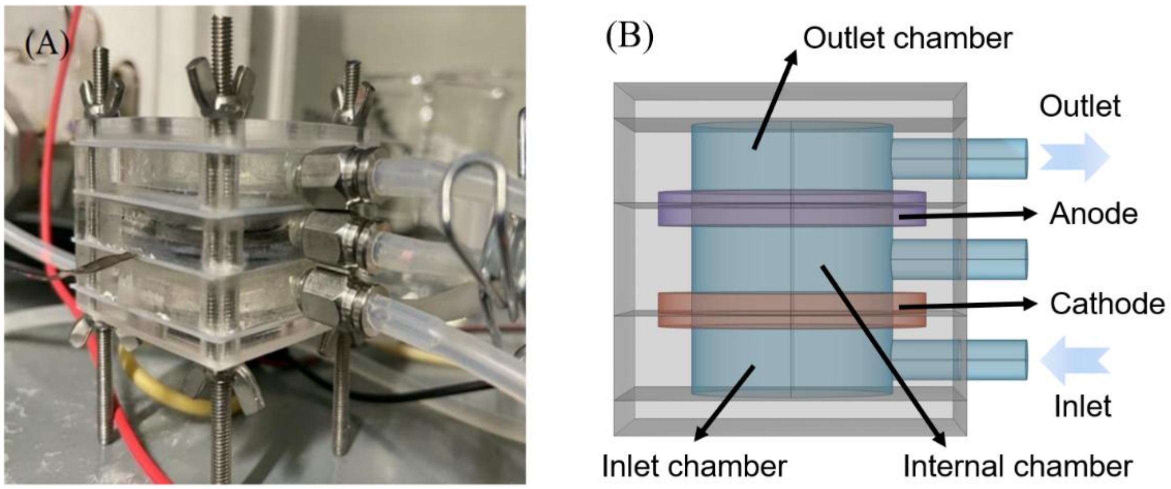

**Figure S2** The (A) photograph and (B) vertically expanded depiction of the flow-through EC reactor.

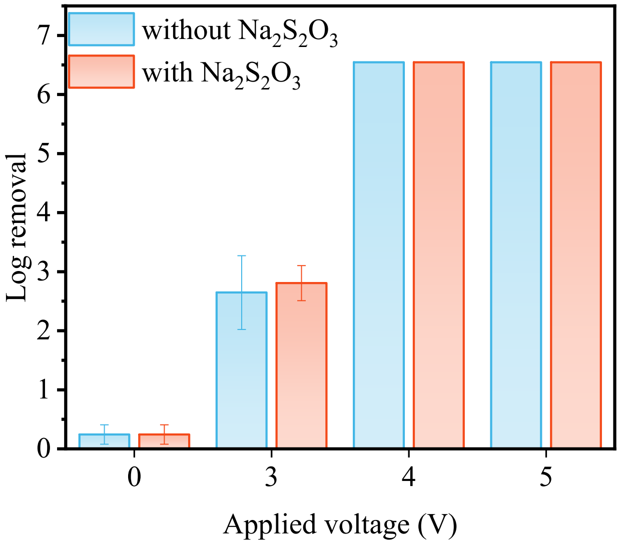

**Figure S3** Verification of the residual disinfecting capacity of the EC reactor effluent.

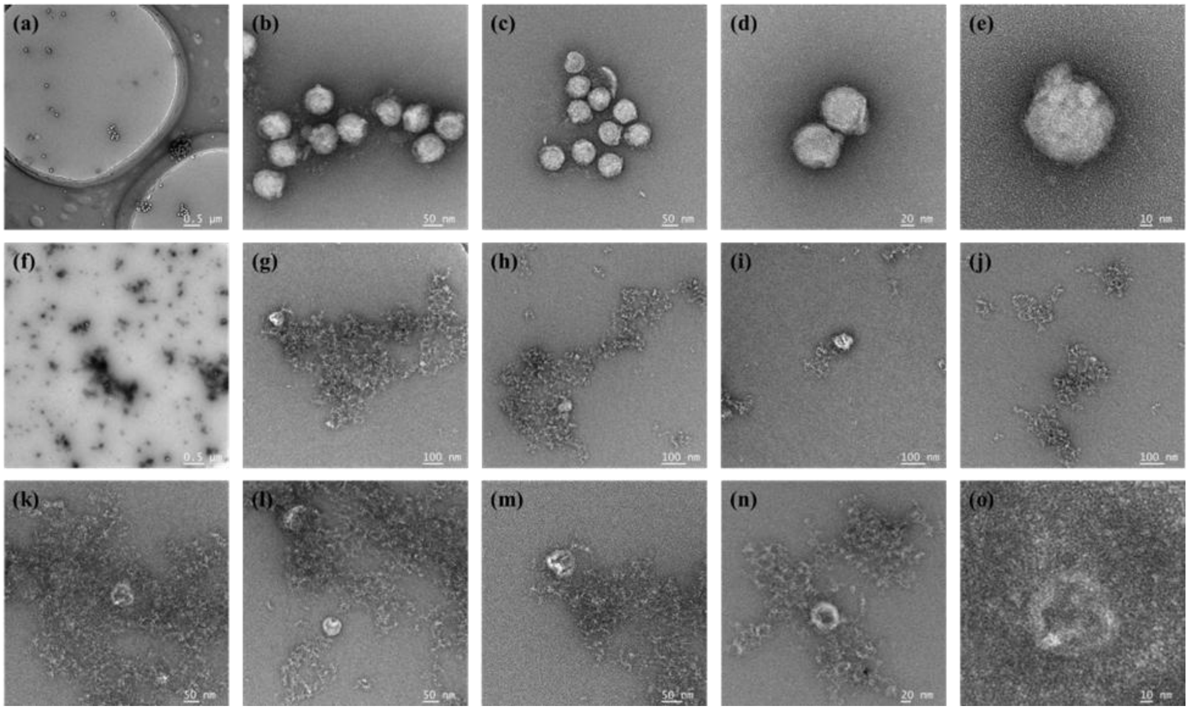

**Figure S4** Full-scale TEM image of Phi6 before (a~e) and after EC disinfection (f~o).

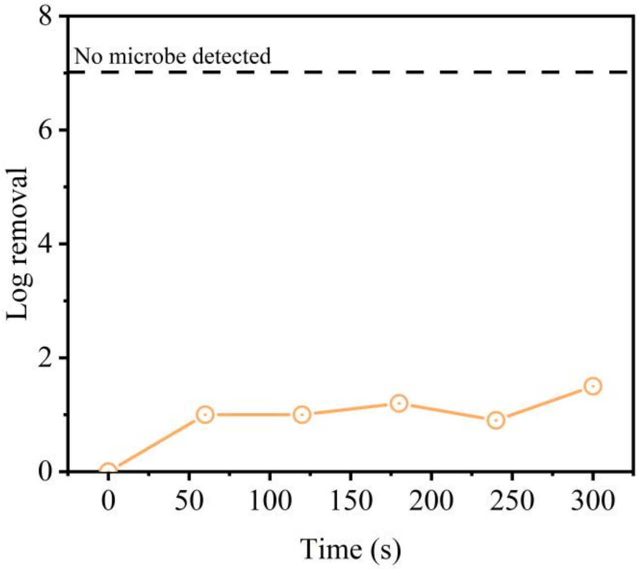

**Figure S5** The time course of Phi6 inactivation efficiency in simulated $H_2O_2$ solution.

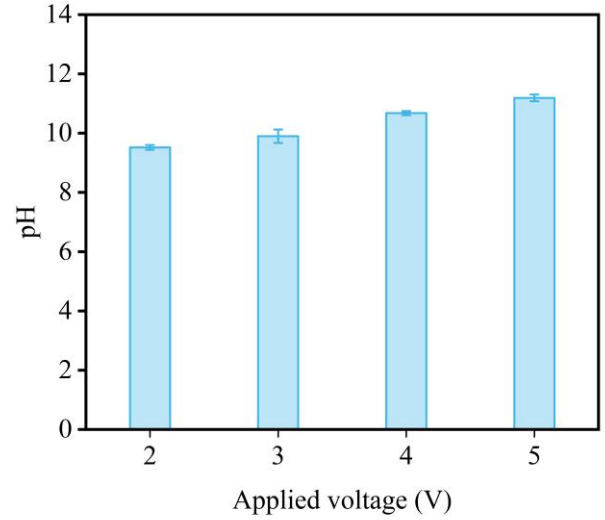

**Figure S6** The change of pH values of cathode effluent with voltages.

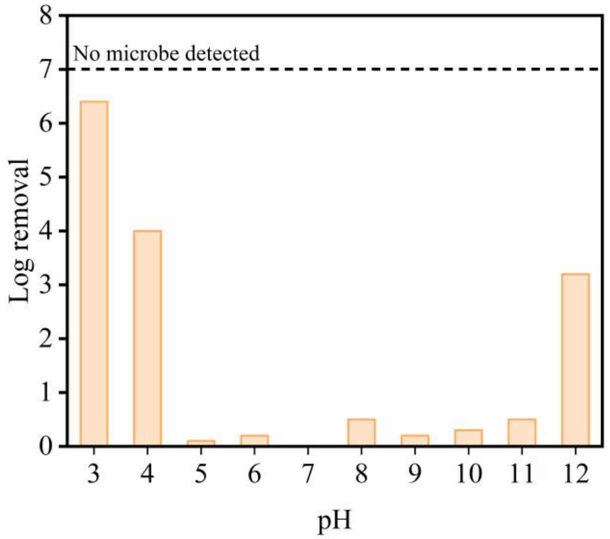

**Figure S7** The activity loss of Phi6 under different pH in 10 mM NaCl solutions.

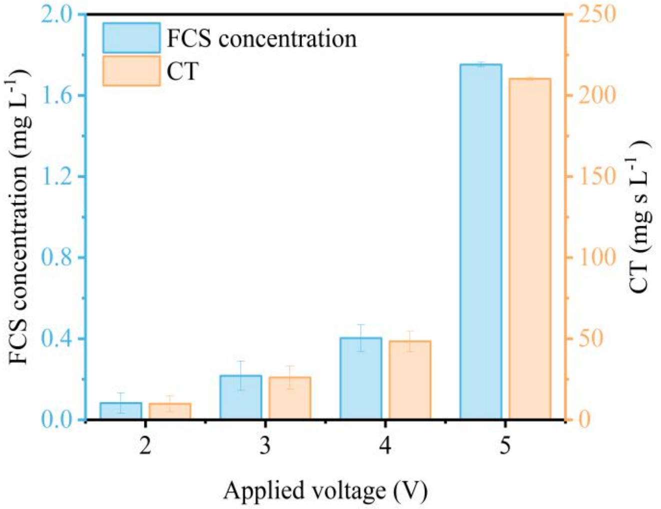

**Figure S8** The FC concentration and the calculated CT value of the EC reactor under different cell voltages.

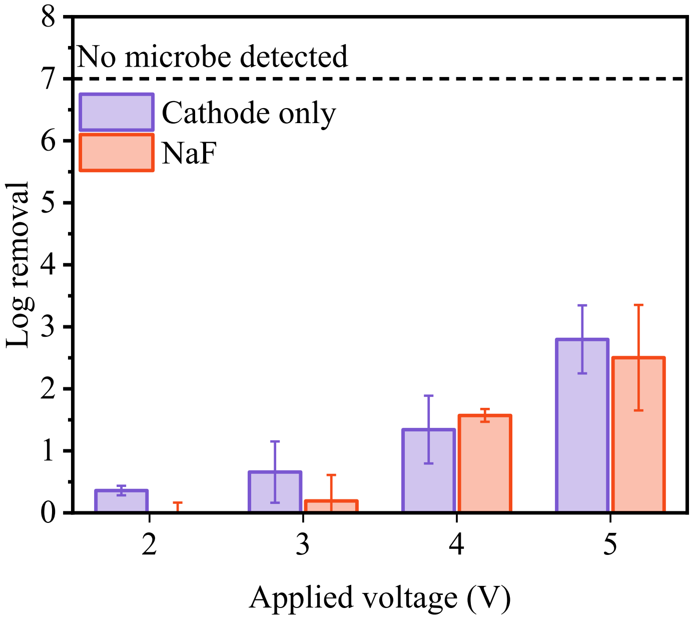

**Figure S9** Contributions of phi6 inactivation to cathodic and anodic EC reactions in 10 mM NaF solution.

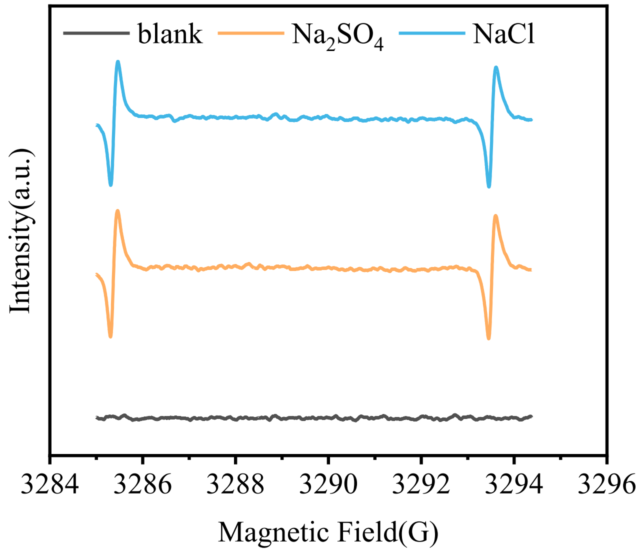

**Figure S10** EPR spectra of DMPO-OH in $Na_2SO_4$ and NaCl solution under 4 V.

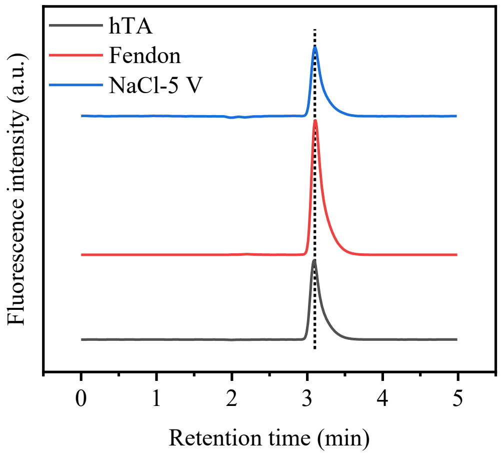

**Figure S11** Comparison of fluorescence signals' retention times in HPLC among the Fenton system, EC reactor effluent, and hTA aqueous solution.

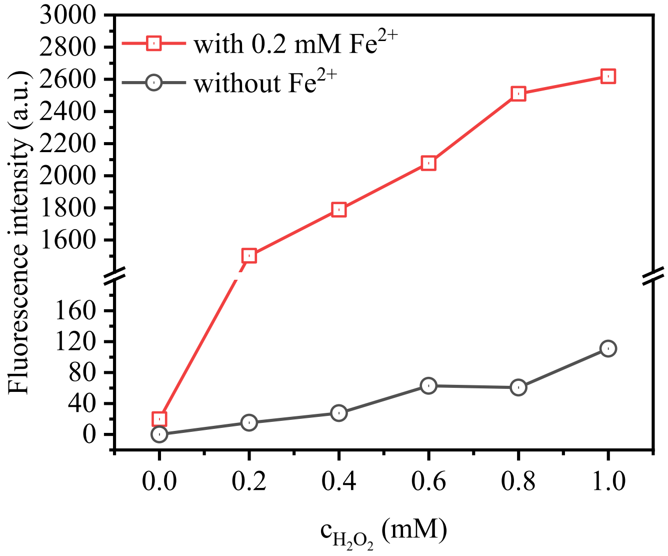

**Figure S12** The Fluorescence intensity production from 100 μM TA in different $H_2O_2$ solutions with and without 0.2 mM $Fe^{2+}$.

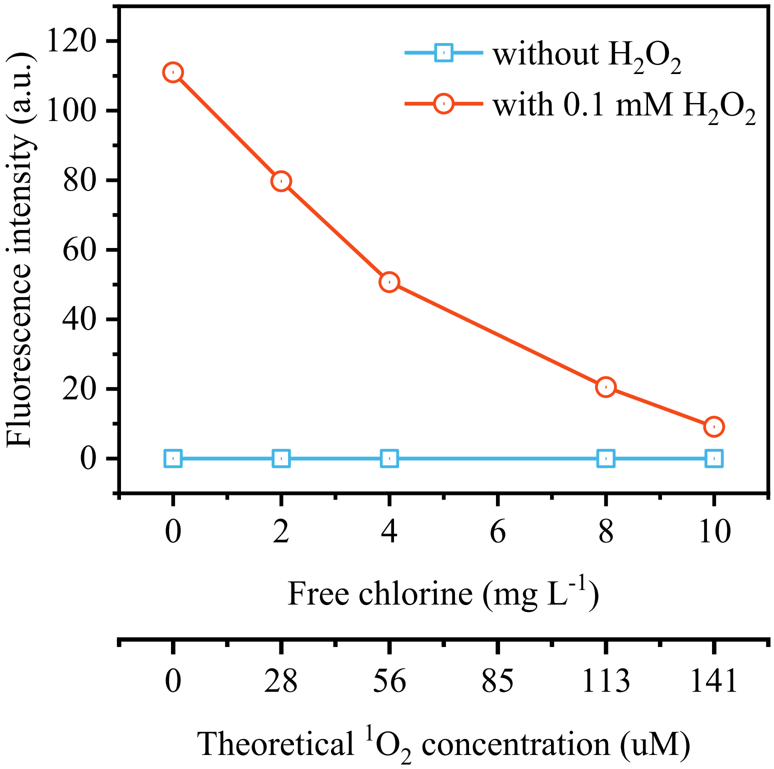

**Figure S13** The FI production from 100 μM TA in different free chlorine solutions with and without 1 mM $H_2O_2$.

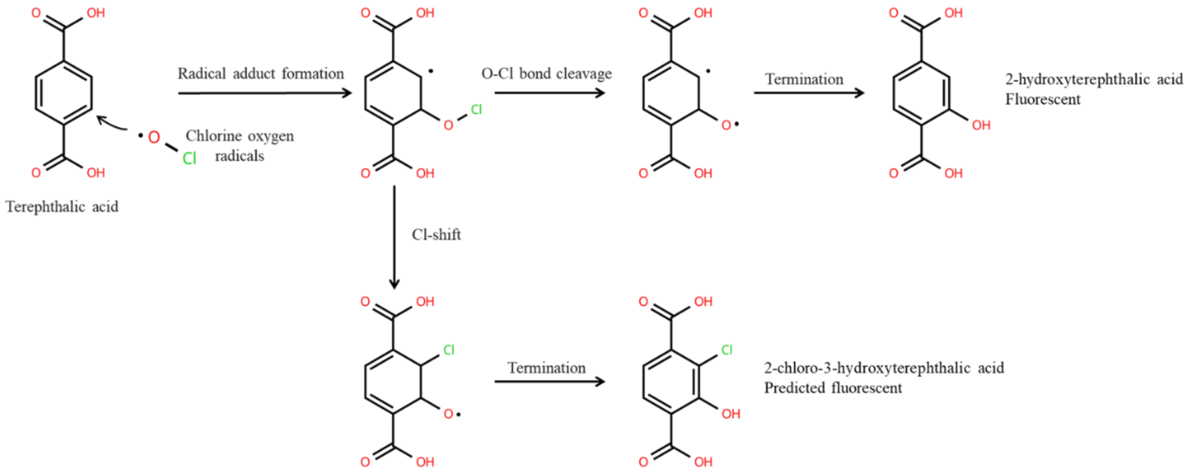

**Figure S14** Possible transformation pathways of TA in the presence of ·ClO.

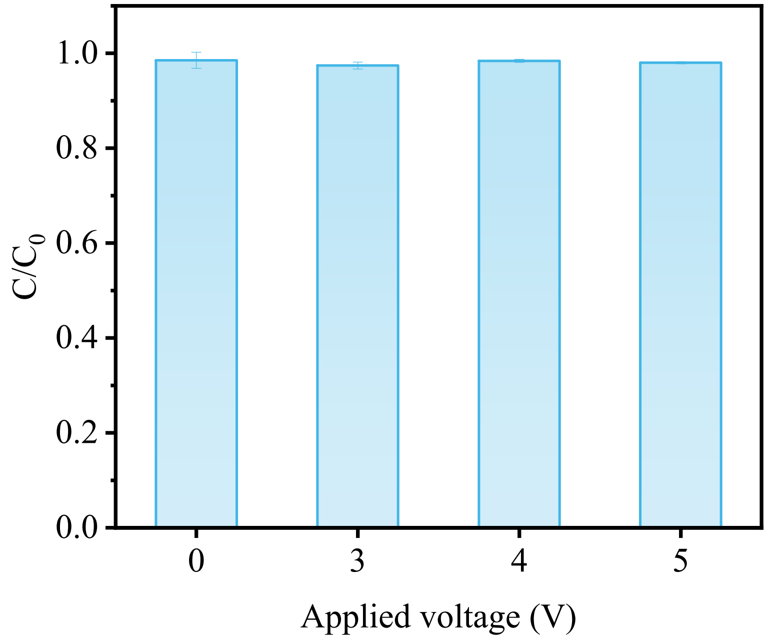

**Figure S15** The probe tests of BA in the flow-through EC reactor in 10 mM NaCl.

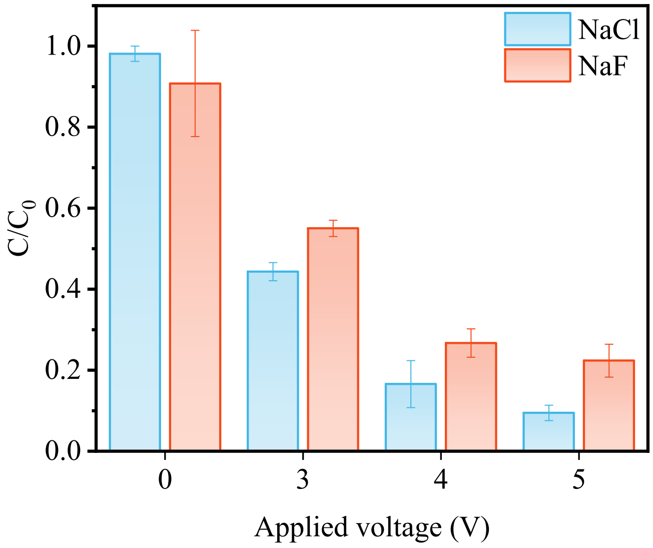

**Figure S16** The probe tests of CBZ in the flow-through EC reactor in 10 mM NaCl and NaF.

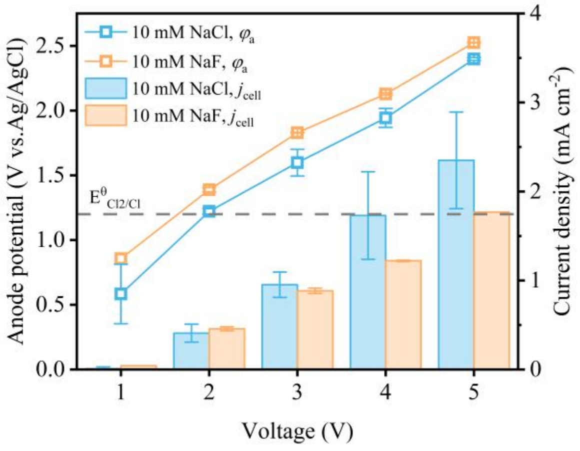

**Figure S17** The anode potential and current under different voltages in the EC reactor.

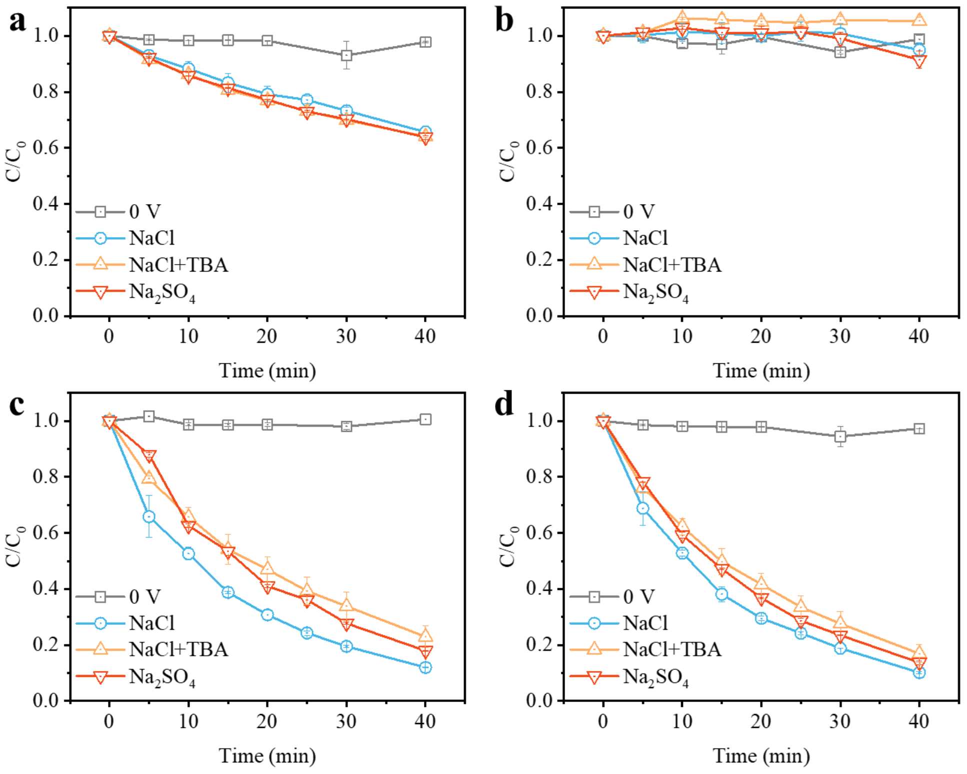

**Figure S18** The kinetics of (a) NB, (b) BA, (c) CBZ, and (d) DMOB in 10 mM NaCl with and without TBA and 5.25 mM $Na_2SO_4$.

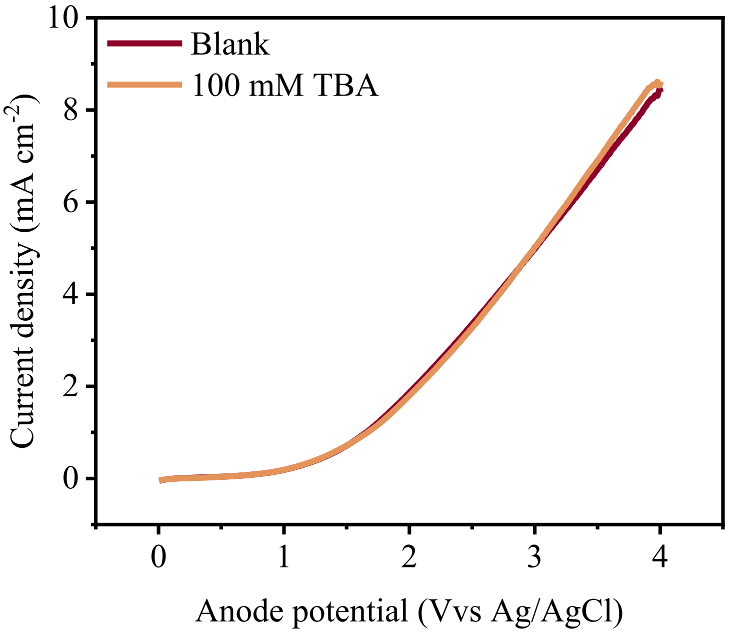

**Figure S19** The LSV curves of CFF in 10 mM NaCl with and without 100 mM TBA.

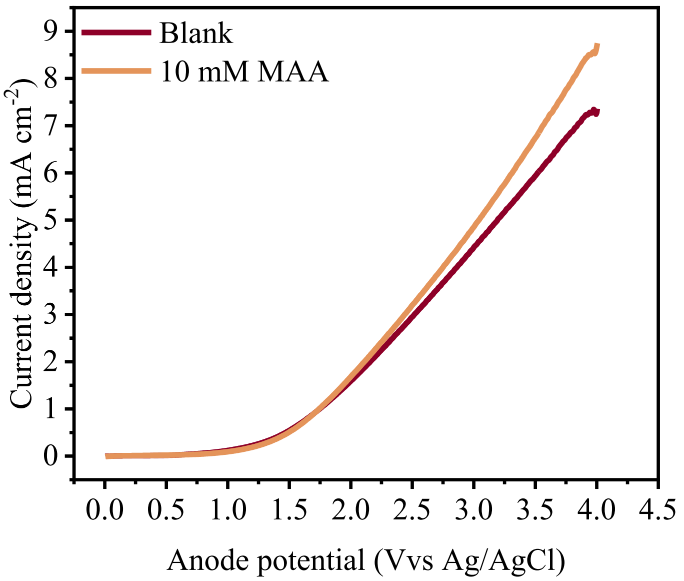

**Figure S20** The LSV curves of CFF in 10 mM NaCl with and without 10 mM MAA.

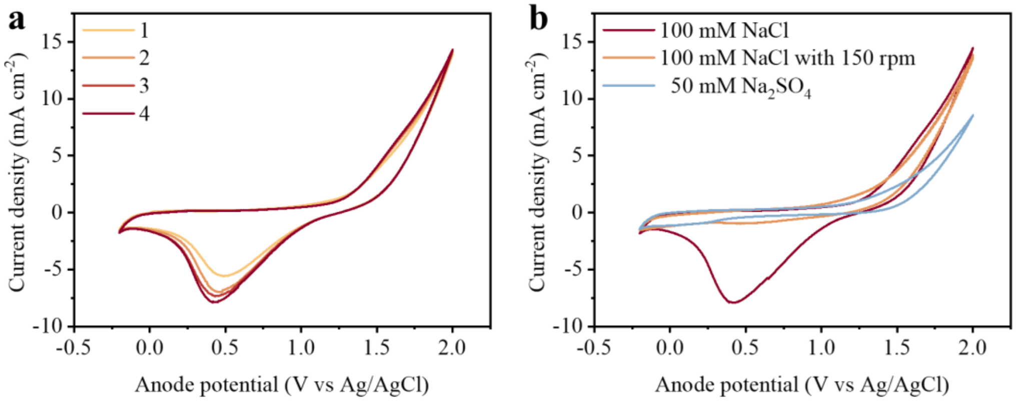

**Figure S21** The CV curves of CFF in (a) 100 mM NaCl and the comparison of the 5th CV curves under different operating conditions as in 50 mM $Na_2SO_4$ and with stirring.

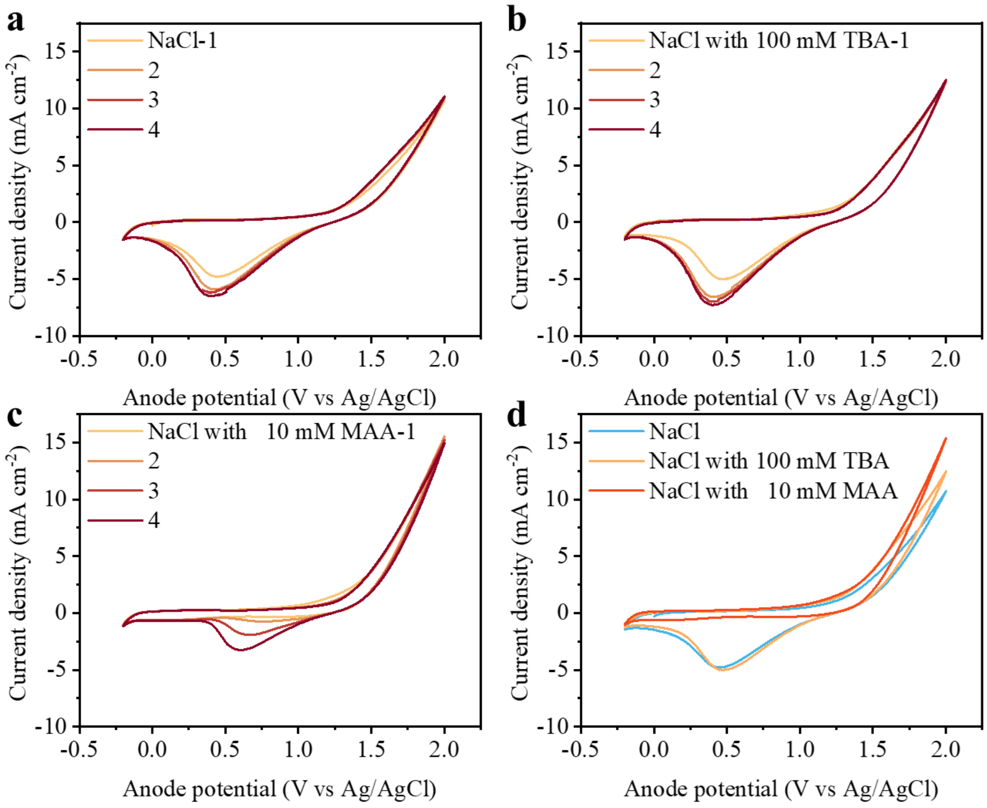

**Figure S22** The CV curves of CFF in(a) 100 mM NaCl, (b) with 100 mM TBA and (c) with 10 mM MAA, and the comparison of 1st curve among different conditions.

**Table**

**Table S1** The physical and chemical properties of CFF.

| Properties | Value |
|---|---|
| Porosity | 90% |
| Conductivity | 380 S $m^{-1}$ |
| Fiber diameter | 9.6 μm |
| Pore diameter | 50-200 μm |
| Specific surface area | 4.5 $m^2$ $g^{-1}$ |

**Table S2** The sequences of amplicon and primer of Phi6 by RT-q-PCR.

| Target | Prime/Amplicon | Sequence |
|---|---|---|
| Phi6 | F | 5’-TGGCGGCGGTCAAGAGC-3’ |
| | R | 5’-GGATGATTCTCCAGAAGCTGCTG-3’ |
| | P | 5’-FAM-CGGTCGTCGCAGGTCTGACACTCGC-3’ |

**Table S3** Verification of PMA staining and photo inactivation.

| Treatment | Fresh Phi6 culture | Extracted Phi6 RNA |
|---|---|---|
| Without PMA staining (CT value for RT-qPCR) | 19.296 ± 0.534 | 22.3887 ± 1.3422 |
| With PMA staining (CT value for RT-qPCR) | 20.2457 ± 0.9452 | N.D.* |

* N.D. means negative results from RT-qPCR

**Table S4** The operation parameters of HPLC analysis for the probe compounds.

| | NB | BA | DMOB | CBZ | TA | hTA |
|---|---|---|---|---|---|---|
| Chromatographic column | C18(5 μm × 10 cm，Agilent) | | | | | |
| Column temperature | 25 ℃ | | | | | |
| Mobile flow rate | 1 mL $min^{-1}$ | | | | | |
| Mobile phase(A) | 0.1% $H_3PO_4$ aqueous solution | | | | | |
| Mobile phase(B) | Acetonitrile | | | | methanol | |
| Mobile phase (A:B, v/v) | 0 min: 50:50; 3 min: 65:35; 4.5 min: 65:35; 5 min: 50:50 | | | | 60:40 | |
| DAD detected wavelength (nm) | 260 | 230 | 211 | 211 | 254 nm | |
| FLD | | | | | $\lambda_{ex}$=315 nm $\lambda_{em}$=425 nm | $\lambda_{ex}$=315 nm $\lambda_{em}$=425 nm |

DAD: Photodiode array detector; FLD: Fluorescence detector; $\lambda_{ex}$: Excitation wavelength; $\lambda_{em}$: Emission wavelength.

**Table S5** Previous research on electrochemical inactivation of viruses.

| **Anode** | ***A***<br>cm$^2$ | ***D***$_i$<br>mm | ***J***<br>mA cm$^{-2}$ | ***E***$_{cell}$<br>V | **Main Process** | **RC** | **Influent** | **Virus** | ***C***$_i$<br>PFU mL$^{-1}$ | ***E***<br>log | **Ref** |
|---|---|---|---|---|---|---|---|---|---|---|---|
| Sno$_2$-Sb REM | 12.6 | 20 | 2.5 | 3.5 | ·OH | FT(AC) | 0.05 M $Na_2SO_4$<br>MWW | MS2 | $10^8$<br>$10^5$ | All | 4 |
| MWNT | 9.6 | | | 2 | EA<br>DO | FT(CA) | 10 mM NaCl | MS2 | $10^6$ | All | 22 |
| MWNT | 9.6 | | | 3 | EA<br>DO | FT(CA) | River water | MS2 | $10^6$ | All | 23 |
| NAT/AT | 6 | | 10 | | ·OH<br>$O_3$ | B | 30 mM $NaClO_4$<br>30 mM NaCl<br>Toilet Wastewater | MS2 | $10^6$ | All | 24 |
| LIG | 16.6 | | 9.2 | 20 | / | FT | PBS | VlV | $10^5$ | All | 25 |
| $BiO_x/TiO_2$ | 90.5 | 5 | 1.2 | 4 | FC | B | Toilet Wastewater<br>with 15 mM NaCl | rAd5<br>MS2 | $10^5$<br>$10^4$ | 3.2<br>All | 5 |
| PPyNW | 0.79 | 10 | / | 1 | EP | FT(CA) | Tap Water | MS2 | $10^4$ | 4 | 6 |
| $Cu_3P$ NW-Cu | 1 | / | 0.107 | 2 | EP, RCS | FT(CA) | 10 mm PBS<br>1.4 mM NaCl | MS2 | $10^7$ | All | 7 |

| **Anode** | ***A*** | ***D*i** | ***J*** | ***E*cell** | **Main Process** | **RC** | **Influent** | **Virus** | ***C*i** | ***E*** | **Ref** |
|---|---|---|---|---|---|---|---|---|---|---|---|
| | $cm^2$ | mm | $mA\ cm^{-2}$ | V | | | | | $PFU\ mL^{-1}$ | log | |
| DSA | 20 | 25 | 3.75 | / | $^1O_2$ | B | 10 mM NaCl<br>2 mM PBS<br>1 mM $H_2O_2$ | MS2 | $10^7$ | >4 | 8 |
| Au | | | | 0.9 | DO | FBC | PBS | FCV | $10^8$ | >5 | 26 |
| $Ti_4O_7$ CM | 7.07 | 5 | 10 | 9.5 | ·OH | FT(AC) | 0.05 M $Na_2SO_4$ | MS2 | $10^{11}$ | 6.7 | 9 |
| LIG-$TiO_x$ | 16.6 | | | 5 | EP<br>FC | FT(CA) | 0.9 % NaCl | MS2<br>Phi6<br>T4 | $10^6$ | 7<br>5.5<br>4 | 10 |
| CFF | 12.6 | 4 | 0.8 | 5 | RaCS | FT(CA) | 10 mM NaCl | Phi6 | $10^7$ | All | |

A: Electrode area; AC: From anode to cathode; B: Batch mode; CA: From cathode to anode; $C_i$: Initial titer; $Cu_3P$ NW-Cu: $Cu(OH)_2$ nanowire-modified Cu mesh; $D_i$: Electrode spacing; DO: Direct oxidation; DSA: Dimensionally stable anode; E: Inactivation efficiency; EA: Electrochemical adsorption; EP: Electroporation; FBC: Cycled flow-by mode; FCV: Feline calicivirus; FT: Flow-through mode; GO–MMO: GO-Modified mixed metal oxide; J: Current density; LIG: Laser-induced graphene; MWNT: Multiwalled carbon nanotube; MWW: Municipal wastewater; NAT/AT: Ni-Sb-$SnO_2$/Sb-$SnO_2$; PBS: Phosphate buffered saline; PPyNW: Polypyrrole nanowire arrays-modified GF electrodes; RC: Reactor configuration; REM: Reactive membrane electrode; $Ti_4O_7$ CM: Magnéli phase $Ti_4O_7$ ceramic membrane; VlV: Vaccinia lister virus.

**Table S6** The basic information of Phi6 protein profiles.

| Protein Number | Molecular Weight (kDa) | Function | Position |
|---|---|---|---|
| P1 | 85.00 | underwear shell structural protein | internal protein |
| P2 | 74.93 | RNA synthetase | internal protein |
| P3 | 68.58 | envelope surface glycoprotein | outer protein |
| P4 | 35.17 | capsid protein | internal protein |
| P5 | 24.03 | invasion of host bacteria and hydrolysis of the host cell wall | membrane protein |
| P6 | 17.36 | unknown | membrane protein |
| P7 | 17.30 | assembling RNA | internal protein |
| P8 | 9.61 | unknown | membrane protein |
| P9 | 7.65 | spinoid protein | membrane protein |

**Table S7** The second-order reaction rate constants between different probes with ·OH, ·Cl, ClO·, and $·Cl_2^-$.

| | Reaction rate($\times 10^9$ $M^{-1}$ $s^{-1}$) | | | |
|---|---|---|---|---|
| | ·OH | ·Cl | ClO· | $·Cl_2^-$ |
| NB | 3.9[1,27] | 0.52[27,28] | <0.0001[27] | 0.0002[27] |
| BA | 5.9[1,27] | 14[1,27] | <0.002[27] | <0.003[27] |
| CBZ | 8.8[1] | 33[1] | 0.197[1] | 0.043[1] |
| DMOB | 7[1,27] | 18[1] | 2.1[1,27] | 0.73[27] |
| TA | 4[6] | - | - | - |
| TBA | 0.59[27] | 0.62[27] | <0.013[27] | $7\times 10^{-7}$ [27] |

**Table S8** The observed removal kinetic constants ($k_{obs}$) of NB, BA, CBZ and DMOB under different conditions.

| Probe | NaCl | | NaCl+TBA | | $Na_2SO_4$ | |
|---|---|---|---|---|---|---|
| | $k_{obs}$ ($10^{-3}$ $min^{-1}$) | $R^2$ | $k_{obs}$ ($10^{-3}$ $min^{-1}$) | $R^2$ | $k_{obs}$ ($10^{-3}$ $min^{-1}$) | $R^2$ |
| NB | 10.8±1.3 | 0.996 | 12.1±0.1 | 0.9932 | 12.1±0.1 | 0.9918 |
| BA | 0.3±0.7 | 0.106 | -1.9±0.6 | 0.8119 | 0.8±0.2 | 0.2089 |
| CBZ | 55.7±1.1 | 0.9953 | 37.6±6.8 | 0.9984 | 42.7±0.2 | 0.9982 |
| DMOB | 57.9±0.9 | 0.9977 | 44.5±7.1 | 0.9991 | 49.6±0.4 | 0.9992 |

**Table S9** The TA transformation by the Fenton system and $H_2O_2$ solution from 100 μM TA.

| $c_{H2O2}$ | with $Fe^{2+}$ | | | without $Fe^{2+}$ | | |
|---|---|---|---|---|---|---|
| | TA removal | FI | $c_{eq,hTA}$ | TA removal | FI | $c_{eq,hTA}$ |
| mM | % | | nM | % | | nM |
| 0 | 2.21 | 19.8 | 21.65 | 0 | 0 | 0 |
| 0.2 | 59.06 | 1502.6 | 1988.37 | 8.14 | 15.1 | 15.09 |
| 0.4 | 66.18 | 1788.7 | 2367.23 | 7.71 | 27.5 | 31.68 |
| 0.6 | 68.82 | 2078.5 | 2749.29 | 8.18 | 62.9 | 79.05 |
| 0.8 | 69.11 | 2509.9 | 3292.05 | 9.16 | 60.7 | 76.1 |
| 1 | 68.49 | 2618.4 | 3487.13 | 8.19 | 111 | 143.4 |

**Table S10** The TA transformation by free chlorine from 100 μM TA.

| $c_{FC}$ (mg $L^{-1}$) | solution | pH | TA removal (%) | | FI production | |
|---|---|---|---|---|---|---|
| | | | 15 min | 30 min | 15 min | 30 min |
| 2 | DI water | 7.15±0.04 | 3.61±0.83 | 3.82±0.7 | 0 | 0 |
| 2 | PBS | 7.3±0.01 | 3.6±0.52 | 3.1±0.48 | 0 | 0 |
| 2 | PBS | 3.24±0.03 | 5.17±2.05 | 6.38±0.3 | 0 | 0 |

**Table S11** The TA transformation by free chlorine and $^1O_2$ from 100 μM TA.

| $c_{H2O2}$ (mM) | $c_{FC}$ (mg $L^{-1}$) | TA removal (%) | | | FI production | | |
|---|---|---|---|---|---|---|---|
| | | 10 min | 20 min | 30 min | 10 min | 20 min | 30 min |
| 0 | 0 | 0 | 0 | 0 | 0 | 0 | 0 |
| 0 | 2 | 1.91 | 1.95 | 1.92 | 0 | 0 | 0 |
| 0 | 4 | 4.5 | 4.46 | 4.45 | 0 | 0 | 0 |
| 0 | 8 | 2.34 | 2.35 | 2.25 | 0 | 0 | 0 |
| 0 | 10 | 3.84 | 3.84 | 3.84 | 0 | 0 | 0 |
| 1 | 0 | 7.46 | 7.77 | 8.19 | 48.1 | 76.8 | 111 |
| 1 | 2 | 8.33 | 8.74 | 8.78 | 22.5 | 53.9 | 79.7 |
| 1 | 4 | 7.73 | 8.17 | 7.91 | 11.6 | 29.7 | 50.7 |
| 1 | 8 | 7.77 | 7.75 | 7.72 | 9.4 | 14.2 | 20.5 |
| 1 | 10 | 7.67 | 7.71 | 7.35 | 6.3 | 6.8 | 9.1 |

**Table S12** Influent parameters, operational parameters, and transformation performance under different electrolyte conditions in TA probe experiments at a voltage of 5 V.

| Electrolyte | Quencher | Conductivity ($\mu S\ cm^{-1}$) | pH | Current density ($mA\ cm^{-2}$) | TA removal (%) | $c_{hTA}$ (nM) |
|---|---|---|---|---|---|---|
| 10 mM NaCl | None | 1250 | 7.82 | 2±0.03 | 4.02±0.6 | 51.58±10.02 |
| 10 mM NaCl | 100 mM TBA | 1223 | 7.69 | 1.98±0.05 | 2.93±1.39 | 0 |
| 10 mM NaCl | 10 mM MAA | 1255 | | 1.95±0.05 | 0.99±0.31 | 0 |
| 10 mM NaF | None | 1047 | 6.67 | 1.7±0.03 | 2.11±0.12 | 0 |
